# MAPK-induced miR-29 targets MAFG and suppresses melanoma development

**DOI:** 10.1101/2020.01.27.922153

**Authors:** Olga Vera, Ilah Bok, Neel Jasani, Koji Nakamura, Xiaonan Xu, Nicol Mecozzi, Ariana Angarita, Kaizhen Wang, Kenneth Y. Tsai, Florian A. Karreth

## Abstract

The tumor suppressive miR-29 family of microRNAs is encoded by two clusters, miR-29b1∼a and miR-29b2∼c, and is regulated by several oncogenic and tumor suppressive stimuli. Here we investigated whether oncogenic MAPK hyperactivation regulates miR-29 abundance and how this signaling axis impacts melanoma development. Using mouse embryonic fibroblasts and human melanocytes, we found that oncogenic MAPK signaling stimulates p53-independent and p53-dependent transcription of pri-miR-29b1∼a and pri-miR-29b2∼c, respectively. Expression analyses revealed that while pri-miR-29a∼bl remains elevated, pri-miR-29b2∼c levels decrease during melanoma progression. Using a rapid mouse modeling platform, we showed that inactivation of miR-29 in vivo accelerates melanoma development and decreases overall survival. We identified the transcription factor MAFG as a *bona fide* miR-29 target that has oncogenic potential in melanocytes and is required for growth of melanoma cells. Our findings suggest that MAPK-induced miR-29 contributes to a tumor suppressive barrier by targeting MAFG, which is overcome by attenuation of miR-29b2∼c expression.

## INTRODUCTION

Malignant melanoma is an aggressive cancer that arises from melanocytes as a consequence of aberrant activation of proto-oncogenes, most commonly BRAF and NRAS. Despite the recent advances in melanoma therapy, most patients still succumb to the disease. Thus, a better understanding of the molecular biology of melanoma is needed to identify new therapeutic targets. In addition to genetic and genomic alterations, aberrant gene expression control contributes to melanoma development and progression. Non-coding RNAs (ncRNAs) are effective regulators of gene expression, especially microRNAs (miRNAs) who repress gene expression by binding to 3’UTRs of target mRNAs (Davis and Hata 2009). To date, various miRNAs have been reported to regulate the biology of melanoma cells (reviewed in (Wozniak et al. 2016; Fattore et al. 2017; Romano and Kwong 2017)) including miR-29a, for which tumor suppressive potential was recently described in vitro (Xiong et al. 2018).

The miR-29 family is encoded by two clusters, miR-29b1∼a and miR-29b2∼c, located on chromosomes 7q32.2 and lq32.2 in humans, respectively (Kriegel et al. 2012; Alizadeh et al. 2019). Expression of the miR-29 clusters produces primary transcripts pri-miR-29b1∼a and pri-miR-29b2∼c, which are processed to generate three mature miRNAs, miR-29a, miR-29b, and miR-29c. The mature miR-29 family members are highly conserved across species and share identical seed sequences (Kriegel et al. 2012). Numerous studies describe the regulation of miR-29 targets, typically focusing on the interaction between target mRNAs and one of the three mature members of the miR-29 family. Given that mature miR-29a, −29b, and −29c share a conserved seed sequence, it is likely that most miR-29 targets have the potential to be repressed by all three miR-29 family members (Kriegel et al. 2012). For instance, SIRT1 has been reported to be regulated by miR-29a in hepatocellular carcinoma and in cervical cancer (Zhang et al. 2018; Nan et al. 2019), miR-29b in colorectal cancer (Liu and Cheng 2018) and miR-29c also in hepatocellular carcinoma (Bae et al. 2014). miR-29 is considered a tumor suppressor miRNA given its ability to repress genes involved in proliferation and cell survival such as AKT3 (Ugalde et al. 2011; Wei et al. 2013), DNMT3A/B (Nguyen et al. 2011), MCL1 (Mott et al. 2007), and CDK6 (Zhao et al. 2010). miR-29 is frequently downregulated in cancer (Alizadeh et al. 2019; Kwon et al. 2019) including leukemia (Havelange et al. 2011), ovarian (Teng et al. 2015), breast (Wang et al. 2017), rhabdomyosarcoma (Muniyappa et al. 2009), gastric (Matsuo et al. 2013) and other cancer types (see (Alizadeh et al. 2019). Over the last decade, miR-29 has emerged as a major regulatory hub that integrates signaling from potent oncogenes and tumor suppressors. Indeed, miR-29 expression is repressed by the oncogenes c-Myc, Hedgehog, and NF-κΒ (Chang et al. 2008; Mott et al. 2010; Zhang et al. 2012). NRF2 was reported to stimulate or suppress miR-29 expression in different cell types (Kurinna et al. 2014; Shah et al. 2015), and p53 promotes miR-29 expression when stimulated by aging or chronic DNA damage (Ugalde et al. 2011). However, the regulation of miR-29 expression and its consequences on melanoma biology are poorly understood.

In this study we examined if oncogenic MAPK signaling represses the expression of the tumor suppressive miR-29 family similar to other oncogenic pathways. Surprisingly, we found that miR-29b1∼a expression is enhanced directly by the MAPK pathway and remains elevated in melanoma. Conversely, MAPK signaling requires p53 to promote miR-29b2∼c expression, which enables the attenuation of miR-29b2∼c through diminished p53 activity during the progression of nevi to frank melanoma. Inactivation of miR-29 in a melanoma mouse model augmented tumor development Finally, we identified MAFG as a miR-29 target whose de-repression may be important for melanoma development.

## RESULTS

### Oncogenic BRAF and KRAS increase the abundance of mature miR-29 in mouse embryonic fibroblasts

We first examined if oncogenic BRAF and KRAS lower miR-29 abundance similar to other oncogenic pathways by ectopically expressing BraF^V600E^ and Kras^G12D^ in wildtype mouse embryonic fibroblasts (MEFs). Both oncogenes enhanced activation of the MAPK pathway but, unexpectedly, provoked an increase of mature miR-29a, −29b, and −29c (Figure 1A). In MEFs, supraphysiological expression of Kras^G12D^ stimulates p53 activity (Tuveson et al. 2004), which could explain the increase in miR-29 levels. As expression of Braf^V600E^ and Kras^G12D^ at physiological levels provokes limited p53 activation, we examined miR-29 expression in MEFs carrying Cre-inducible endogenous Braf^V600E^ or Kras^G12D^ alleles (LSL-Braf^V600E^ and LSL-Kras^G12D^) (Jackson et al. 2001; Perna et al. 2015). Adenoviral-Cre mediated activation of endogenous Braf^V600E^ or Kras^G12D^ increased miR-29a, −29b, and −29c levels (Figure 1B and Supplementary Figure 1A), indicating that endogenous expression of these oncogenes also enhances the expression of the miR-29 family. To determine if the regulation of miR-29 occurs at the transcriptional level, we assessed the expression of the primary pri-miR-29b1∼a and pri-miR-29b2∼c transcripts by qRT-PCR. Ectopic overexpression or endogenous activation of oncogenic Braf^V600E^ or Kras^G12D^ increased both pri-miR-29b1∼a and pri-miR-29b2∼c (Figure 1C, 1D and Supplementary Figure 1B), indicating that oncogenic BRAF and KRAS promote the transcription of both miR-29 clusters.

**Figure 1:**
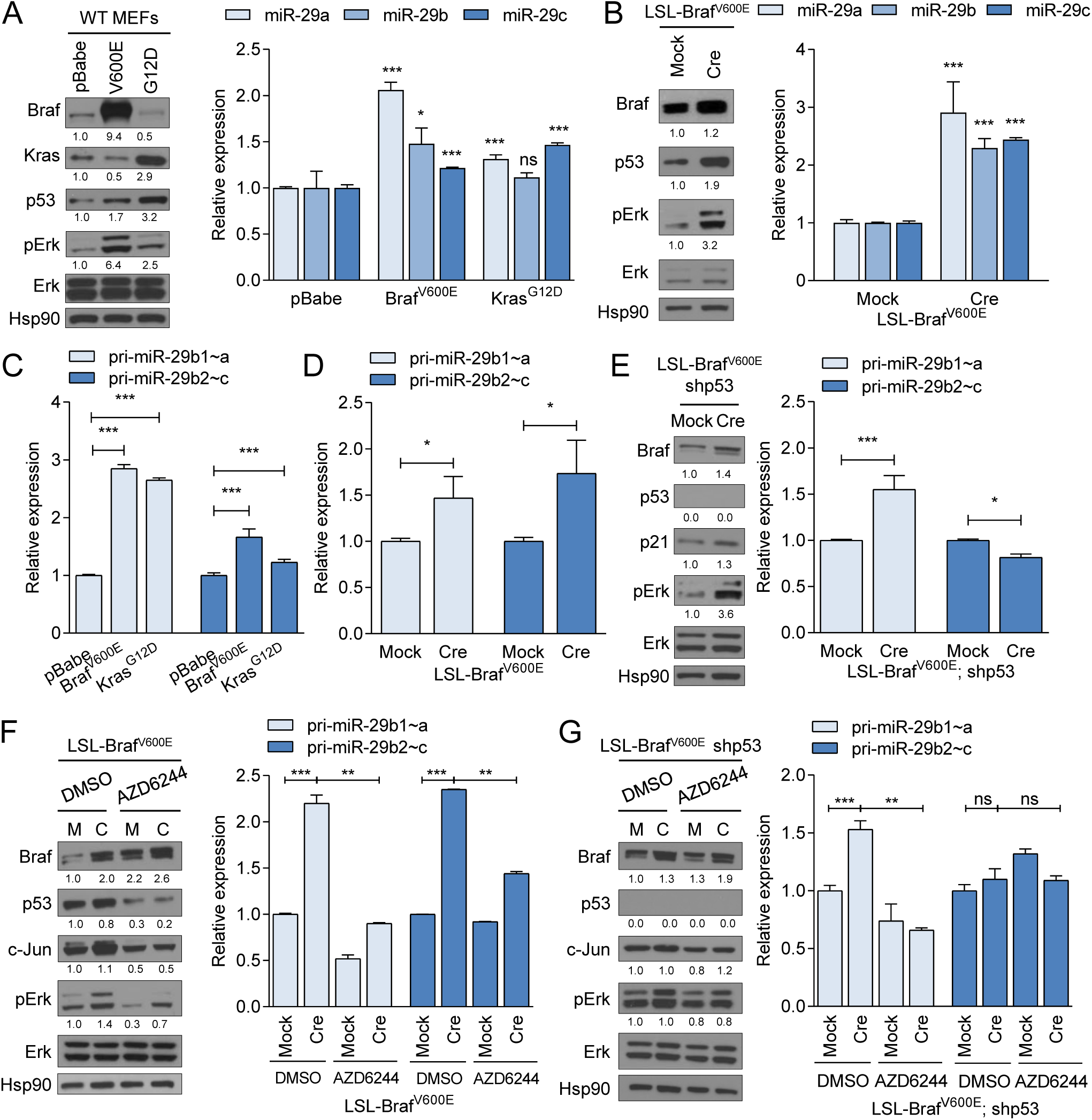
Oncogenic BRAF and KRAS regulate miR-29 in MEFs. (A) Ectopic expression of Braf^V600E^ or Kras^G12D^ in wildtype MEFs elevates expression of p53 and pERK (Western blot, left panel) and mature miR-29a, miR-29b, and miR-29c (qRT-PCR, right panel). (B) Induction of Braf^V600E^ expression by Adeno-Cre (Cre) in LSL-Braf^V600E^ MEFs elevates expression of p53 and pERK (Western blot, left panel) and mature miR-29a, miR-29b, and miR-29c (qRT-PCR, right panel). MEFs infected with empty adenovirus (Mock) serve as controls. (C-D) qRT-PCRs showing the expression of pri-miR-29b1∼a and pri-miR-29b2∼c in wildtype MEFs following overexpression of Braf^V600E^ or Kras^G12D^ (C) and in LSL-Braf^V600E^ MEFs following Adeno-Cre/Mock infection (D). (E) Effect of p53 silencing on pri-miR-29b1∼a and pri-miR-29b2∼c expression measured by qRT-PCR in LSL-Braf^V600E^ MEFs after induction of Braf^V600E^ with Adeno-Cre. (F-G) Effect of MEK inhibitor (AZD6244) on the MAPK pathway and pri-miR-29b1∼a and pri-miR-29b2∼c expression in LSL-Braf^V600E^ (F) and shp53-expressing LSL-Braf^V600E^ MEFs (G). Western blots are shown in the left panels and expression of pri-miR-29b1∼a and pri-miR-29b2∼c measured by qRT-PCR are shown in the right panels of (E), (F) and (G). For all Western blots, Hsp90 was used as loading control. The mean ± SEM of one representative out of two independent experiments performed in triplicate in three different cell lines is shown. Gene expression levels are normalized to U6 (mature miR-29) or β-Actin (pri-miR-29) as endogenous control, ns, not significant; * p < 0.05; ** p < 0.01; *** p < 0.001.

### pri-miR-29b1∼a and pri-miR-29b2∼c are transcribed via the MAPK pathway and p53

To further assess if p53 plays a role in the regulation of miR-29 by oncogenic BRAF and KRAS, we first ascertained that p53 activation enhances the expression of miR-29 in MEFs, as measured by mature miRNA qRT-PCR (Supplementary Figure 1C and 1D). We also tested if induction of p53 increased miR-29 transcription and found that Doxorubicin and Mitomycin C induced only the transcription of pri-miR-29b2∼c (Supplementary Figure 1E). The increase of miR-29a is likely explained by the inability of the mature miRNA qRT-PCR assay to distinguish between miR-29a and miR-29c, which only differ in one nucleotide, as has been suggested previously (Kurinna et al. 2014). Indeed, miR-29a and miR-29c Taqman qRT-PCR assays were unable to distinguish between miR-29a and miR-29c mimics transfected into A375 melanoma cells (Supplementary Figure 1F). Thus, p53 regulates expression of only miR-29b2∼c, while oncogenic BRAF and KRAS induce transcription of both miR-29b1∼a and miR-29b2∼c.

To further test whether Braf^V600E^-induced pri-miR-29b1∼a expression is independent of p53 we silenced p53 in LSL-Braf^V600E^ MEFs (Supplementary Figure 1G). Retroviral shRNA infection did not affect Braf^V600E^-mediated pri-miR-29b1∼a and pri-miR-29b2∼c induction (Supplementary Figure 1H). Expression of Braf^V600E^ failed to increase pri-miR-29b2∼c expression in the absence of p53 (Figure 1E) while the induction of pri-miR-29b1∼a was unaffected (Figure 1E), indicating that Braf^V600E^-mediated regulation of pri-miR-29b1∼a in MEFs is indeed independent of p53. To test the functional consequence of Braf^V600E^-induced pri-miR-29b1∼a expression, we performed a miR-29 Luciferase reporter assay in p53-silenced LSL-Braf^V600E^ MEFs. Induction of Braf^V600E^ decreased

Luciferase activity (Supplementary Figure 1I), suggesting that the extent of miR-29b1∼a upregulation is sufficient for target repression.

Given that Braf^V600E^-induced miR-29b1∼a expression is independent of p53, we examined the involvement of the MAPK pathway in miR-29 regulation. To this end, we treated LSL-Braf^V600E^ MEFs with the MEK inhibitor AZD6244 (Selumetinib) for 24 hours. AZD6244 treatment decreased basal and BraF^600E^-induced expression of pErk and the MAPK pathway transcriptional target c-Jun in the presence or absence of p53 (Figure 1F and 1G). pri-miR-29b1∼a expression was similarly reduced by MEK inhibition (Figure 1F and 1G), indicating that oncogenic BRAF regulates miR-29b1∼a independently of p53 via the MAPK pathway. AZD6244 treatment blunted Braf^V600E^-induced pri-miR-29b2∼c expression, possibly due to reduced p53 expression (Figure 1F). Neither Braf^V600E^ expression nor MEK inhibition affected pri-miR-29b2∼c levels in the absence of p53 (Figure 1G). These data suggest differential regulation of the miR-29 clusters by Braf^V600E^: while miR-29b1∼a is controlled by the MAPK pathway, miR-29b2∼c expression depends on p53.

### The MAPK pathway regulates miR-29 expression in melanocytes and melanoma cells

We next examined if the regulation of miR-29 observed in MEFs is conserved in melanocytes. Induction of Braf^V600E^ by adenoviral Cre in LSL-Braf^V600E^ primary mouse melanocytes increased the expression of both miR-29 clusters (Figure 2A)., suggesting that the regulation of miR-29 by oncogenic BRAF is conserved in mouse and human melanocytes. Additionally, treatment of the immortalized human melanocyte lines Hermes1 and Hermes3A with Doxorubicin increased p53, p21, and pri-miR-29b2∼c (Supplementary Figure 2A), indicating that p53 regulates pri-miR-29b2∼c also in melanocytes. To analyze miR-29 regulation by the MAPK pathway in melanocytes we starved Hermes1 and Hermes3A cells of TP A, a phorbol ester that stimulates the MAPK pathway by activating PKC (Schonwasser et al. 1998) and that is required for melanocyte proliferation *in vitro*. Re-stimulation with TPA increased pERK levels and transcription of c-JUN, pri-miR-29b1∼a, and pri-miR-29b2∼c (Figure 2B, Supplementary Figure 2B). Further, MEK inhibition in Hermes1 and Hermes3A cells cultured in the presence of TPA diminished the expression of pri-miR-29b1∼a and pri-miR-29b2∼c (Figure 2C). AZD6244 also decreased the levels of pri-miR-29b1∼a in a panel of BRAF and NRAS mutant human melanoma cell lines, while pri-miR-29b2∼c was only moderately reduced in three out of eight cell lines (Figure 2D and 2E). A similar effect was observed in BRAF mutant lung and colon cancer cells (Supplementary Figure 2C), suggesting that MAPK-mediated regulation of miR-29 is not limited to cells of the melanocytic lineage.

**Figure 2:**
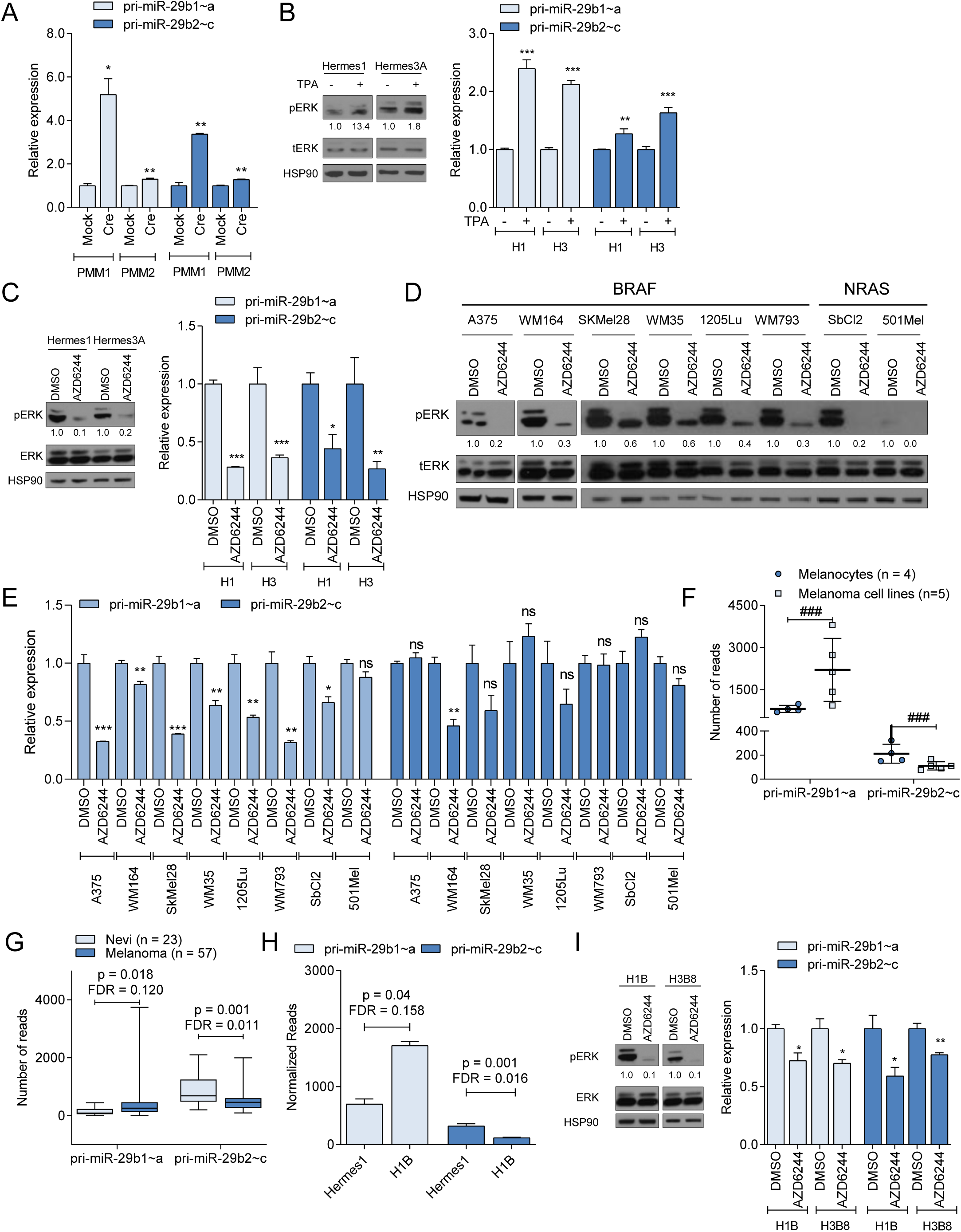
MAPK signaling and p53 regulate miR-29 in human melanocytes and melanoma. (A) qRT-PCRs showing the expression of pri-miR-29b1∼a and pri-miR-29b2∼c in LSL-Braf^V600E^ primary mouse melanocyte (PMM) following Adeno-Cre/Mock infection (A). (B) Effect of TPA on pri-miR-29b1∼a and pri-miR-29b2∼c expression in human melanocytes. (D-F) Effect of AZD6244 on pri-miR-29b1∼a and pri-miR-29b2∼c expression in human melanocytes (D) and BRAF and NRAS mutant melanoma cells (D and E). (F) Expression of pri-miR-29b1∼a and pri-miR-29b2∼c in human melanocytes (n = 4) and melanoma cell lines (n = 5) obtained by RNAseq. (G) Expression of pri-miR-29b1∼a and pri-miR-29b2∼c in nevi (n = 23) and melanoma (n = 57) in the GSE112509 dataset (H) Expression of pri-miR-29b1∼a and pri-miR-29b2∼c in parental Hermes1 and H1B cells obtained by RNAseq. (I) Effect of AZD6244 on pri-miR-29b1∼a and pri-miR-29b2∼c expression in H1B and H3B8 cells. pri-miR-29b1∼a and pri-miR-29b2∼c qRT-PCRs are shown in the right panels while Western blots are shown in the left panels of (C) and (I). For all Western blots, HSP90 was used as loading control. The mean ± SEM of one representative out of two independent experiments performed in triplicate is shown. Gene expression levels are normalized to β-Actin as endogenous control, ns, not significant; * p < 0.05; ** p < 0.01; *** p < 0.001; # FDR < 0.05; ## FDR < 0.01; ### FDR < 0.001.

Our results indicate that Braf^V600E^-induced expression of miR-29 may contribute to a tumor suppressive barrier that protects melanocytes from cellular transformation. To analyze pri-miR-29 expression during melanomagenesis, we performed RNA sequencing of four wildtype BRAF human melanocyte cell lines (Hermes1, Hermes2, Hermes3A, and Hermes4B, immortalized with TERT and CDK4^R24C^ or pl6 silencing) and five human melanoma cell lines (WM35, 1205Lu, WM164, SKMel28 and WM793) harboring the oncogenic Braf^V600E^ mutation. In addition, we interrogated pri-miR-29 expression in a publicly available RNAseq dataset (Kunz et al. 2018) containing 23 nevi and 57 primary melanomas. We found a trend towards increased pri-miR-29b1∼a expression in melanoma cell lines and primary melanomas compared to melanocyte cell lines and nevi, respectively (Figure 2F and 2G). Notably, expression of pri-miR-29b2∼c was attenuated in melanoma cell lines and primary melanomas (Figure 2F and 2G), suggesting that reduced expression of the miR-29b2∼c cluster is associated with melanoma progression. Impaired activity of p53 could cause reduced miR-29b2∼c expression, and we indeed observed diminished p53 and/or p21 induction upon Doxorubicin treatment in melanoma cells compared to melanocytes (Supplementary Figure 2Aand 2D).

To further analyze miR-29 regulation during melanocyte transformation, we examined the consequences of chronic Braf^V600E^ expression in human melanocytes. We delivered lentiviral HA-tagged Braf^V600E^ to Hermes1 and Hermes3A cells, which resulted in the emergence of four independent clones, one from Hermes1 (H1B) and three from Hermes3A (H3B2, H3B4 and H3B8). These cell lines are morphologically different from the parental lines, express ectopic Braf^V600E^, and exhibit increased MAPK signaling as shown by elevated pERK and c-JUN (Supplementary Figure 2E and 2F), which enables these cells to proliferate in the absence of TPA (Supplementary Figure 2G and 2H). Notably, all four cell lines lost expression of p53 (Supplementary Figure 2E and 2F), and RNA sequencing revealed that similar to melanoma cell lines and primary melanomas, H1B cells exhibited reduced pri-miR-29b2∼c expression and a trend towards increased pri-miR-29b1∼a levels (Figure 2H). Finally, expression of both pri-miR-29b1∼a and pri-miR-29b2∼c was sensitive to AZD6244 treatment in H1B and H3B8 cells (Figure 2l). These observations indicate that while MAPK signaling enhances transcription of miR-29b1∼a in melanocytes, miR-29b2∼c is regulated by the MAPK pathway and p53. Furthermore, progression of nevi to frank melanoma is associated with decreased pri-miR-29b2∼c levels, possibly through impaired p53 activity.

### miR-29 inactivation promotes melanoma formation

To examine the role of miR-29 in melanoma formation *in vivo*, we first tested a miRNA sponge approach to inactivate miR-29 *in vitro*. To this end, we either transfected A375 melanoma cells with a hairpin inhibitor of miR-29a or delivered a lentiviral bulged miR-29 sponge construct The miR-29 sponge enhanced the focus formation capacity of A375 cells similar to the miR-29 hairpin inhibitor (Supplementary Figure 3A and 3B). In addition, the miR-29 sponge increased the activity of a miR-29 Luciferase reporter (Supplementary Figure 3C). Given the extensive overlap of predicted miR-29 targets in human and mouse, we used this miR-29 sponge construct in combination with our melanoma mouse modeling platform (Bok et al. 2019) to assess the effect of miR-29 inactivation in melanoma development Embryonic stem cell (ESC)-derived Braf^V600E^; Pten^FL^/wT chimeras produced by this approach are topically treated with 40H-Tamoxifen (4-OHT) to activate melanocyte-specific Tyr-CreERt2, which induces BraF^600E^ expression and heterozygous Pten deletion (Supplementary Figure 3D), thereby initiating melanomagenesis. Cre also induces reverse transactivator (rtTA3) expression, enabling melanocyte-specific expression of transgenes upon Doxycycline administration (Supplementary Figure 3D). We targeted ESCs with a Doxycycline (Dox)-inducible, GFP-linked miR-29 sponge allele or GFP as a control and produced miR-29 sponge and GFP control chimeras. Notably, chimeras expressing the miR-29 sponge developed melanoma with shorter latency (Figure 3A) and exhibited reduced overall survival (Figure 3B), indicating that inactivation of miR-29 accelerates melanoma development. Moreover, while all control mice developed only one melanoma, 37.5% of miR-29 sponge mice developed more than one tumor (Figure 3C). All tumors expressed the melanoma marker S100, confirming their melanocytic origin (Figure 3D).

**Figure 3:**
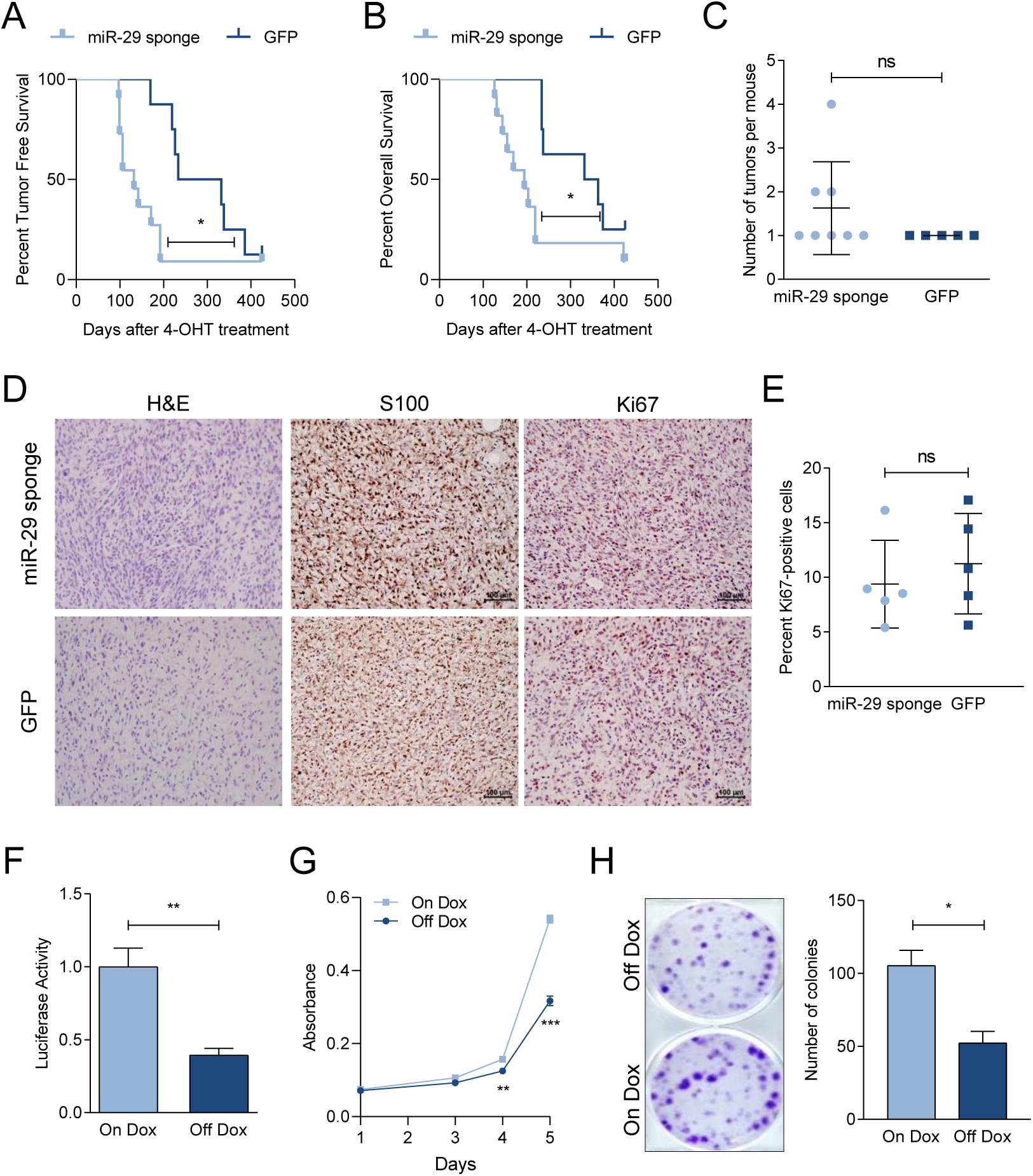
miR-29 inactivation promotes melanoma formation. (A, B) Kaplan-Meier curves comparing the tumor free survival (A) and overall survival (B) of Braf^V600E^; Pten^Δ/WT^ GFP (n = 6) and miR-29 sponge (n = 10) chimeras using Gehan-Breslow-Wilcoxon test. (C) Number of melanomas that developed in the chimeras. (D) Hematoxilin-Eosin and S100 and Ki67 immunohistochemistry staining on tumors from GFP and miR-29 sponge mice at endpoint. (E) Quantification of Ki67-positive nuclei per field in tumors from GFP and miR-29 sponge mice. (F) miR-29-Luciferase reporter activity in miR-29 sponge melanoma cells. Dox withdrawal turns off expression of the sponge construct resulting in miR-29 reactivation. The combined mean ± SEM of three independent experiments performed in quadruplicate is shown. (G, H) Proliferation (G) and colony formation (H) upon miR-29 reactivation in miR-29 sponge melanoma cells. The mean ± SEM of one representative out of three independent experiments performed in quadruplicate is shown, ns, not significant; * p < 0.05; ** p < 0.01; *** p < 0.001

Histologically, GFP control tumors are characterized by small spindled cells often with loose edematous stroma while miR-29 sponge tumors are more punctuated by hypercellular areas with readily-appreciated mitoses. However, immunostaining of Ki67 revealed no difference in melanoma cell proliferation (Figure 3D and 3E), suggesting that miR-29 inactivation has more pronounced effects on tumor initiation or the transition from nevi to melanoma than on progression. Accordingly, we did not observe any gross metastases in miR-29 sponge or GFP control mice.

### Melanoma cells are addicted to high levels of miR-29 target genes

We derived a melanoma cell line from a miR-29 sponge chimera to validate the functionality of the miR-29 sponge in mice and to examine the effects of restoring miR-29 activity. Withdrawing Dox from the melanoma cell line turned off miR-29 sponge expression and enhanced repression of a miR-29 Luciferase reporter (Figure 3F), confirming that the sponge inactivated endogenous miR-29. Moreover, Dox withdrawal reduced proliferation (Figure 3G) and colony formation (Figure 3H) of the miR-29 sponge melanoma cells, indicating that continued miR-29 inactivation, and thus overexpression of miR-29 targets, supports the transformed state.

miR-29 may elicit its tumor suppressive potential by repressing targets such as AKT3, DNMT3A/B, or MCL1 (Mott et al. 2007; Nguyen et al. 2011; Ugalde et al. 2011; Wei et al. 2013); however, miR-29 hairpin inhibitors failed to increase the expression of these validated targets in A375 melanoma cells (Supplementary Figure 4A). Thus, alternative miR-29 targets must be responsible for the observed phenotypes in human cells and our miR-29 sponge melanoma mouse model (Figure 3, Supplementary Figure 3). To identify miR-29 targets with relevance to melanoma, we transfected A375 cells with miR-29a mimics and performed RNA sequencing. 6,358 genes were differentially expressed in response to miR-29a mimics, and of those 1,309 were downregulated with a Log2FC < −0.5 and a FDR < 0.05. We further prioritized potential miR-29 targets based on three criteria: 1) the presence of high-confidence conserved miR-29 binding sites predicted by at least eight different bioinformatics algorithms, 2) increased expression in primary melanoma compared to nevi in the GSE112509 dataset with a Log2FC ≥ 0.3 and a FDR ≤ 0.05, and 3) a negative correlation in expression with pri-miR-29b2∼c in the GSE112509 dataset with r ≤ −0.3 (Supplementary Figure 4B). This analysis yielded 9 candidate target genes: KCTD5, MYBL2, SLC31A1, MAFG, RCC2, TUBB2A, SH3BP5L, SMS, and NCKAP5L (Supplementary Figure 4C and 4D).

### MAFG and MYBL2 are putative miR-29 targets with roles in melanoma

Analyzing the TCGA-SKCM cohort revealed that high expression of the 9 identified candidate miR-29 targets is associated with poorer survival of melanoma patients (Figure 4A), suggesting oncogenic roles for these miR-29 targets in melanoma. To reveal the target(s) affecting melanoma biology, we first tested if miR-29 regulated their expression. Restoring miR-29 activity by withdrawing Dox from the murine miR-29 sponge melanoma cell line increased the expression of 8 out of the 9 identified targets (Figure 4B). Moreover, miR-29 miRNA mimics decreased expression of the 9 targets, while inhibition of endogenous miR-29 with hairpin inhibitors increased expression of 7 out of 9 targets in the murine miR-29 sponge melanoma cells (Figure 4C) and human A375 cells (Figure 4D). Next, to test if repression by miR-29 impairs melanoma cell growth, we silenced the 9 miR-29 targets using individual siRNA pools. Knockdown of *MAFG* and, to a lesser extent, *MYBL2* significantly decreased proliferation (Figure 4E) and focus formation ability (Figure 4G and 4F) of A3 75 melanoma cells. MAFG has been described as an epigenetic regulator and transcriptional repressor in melanoma and hyperactive MAPK may increase MAFG stability by ERK-mediated phosphorylation (Fang et al. 2016). Given these observations and the significant effect of MAFG silencing on melanoma cell growth, we selected MAFG for further analysis.

**Figure 4:**
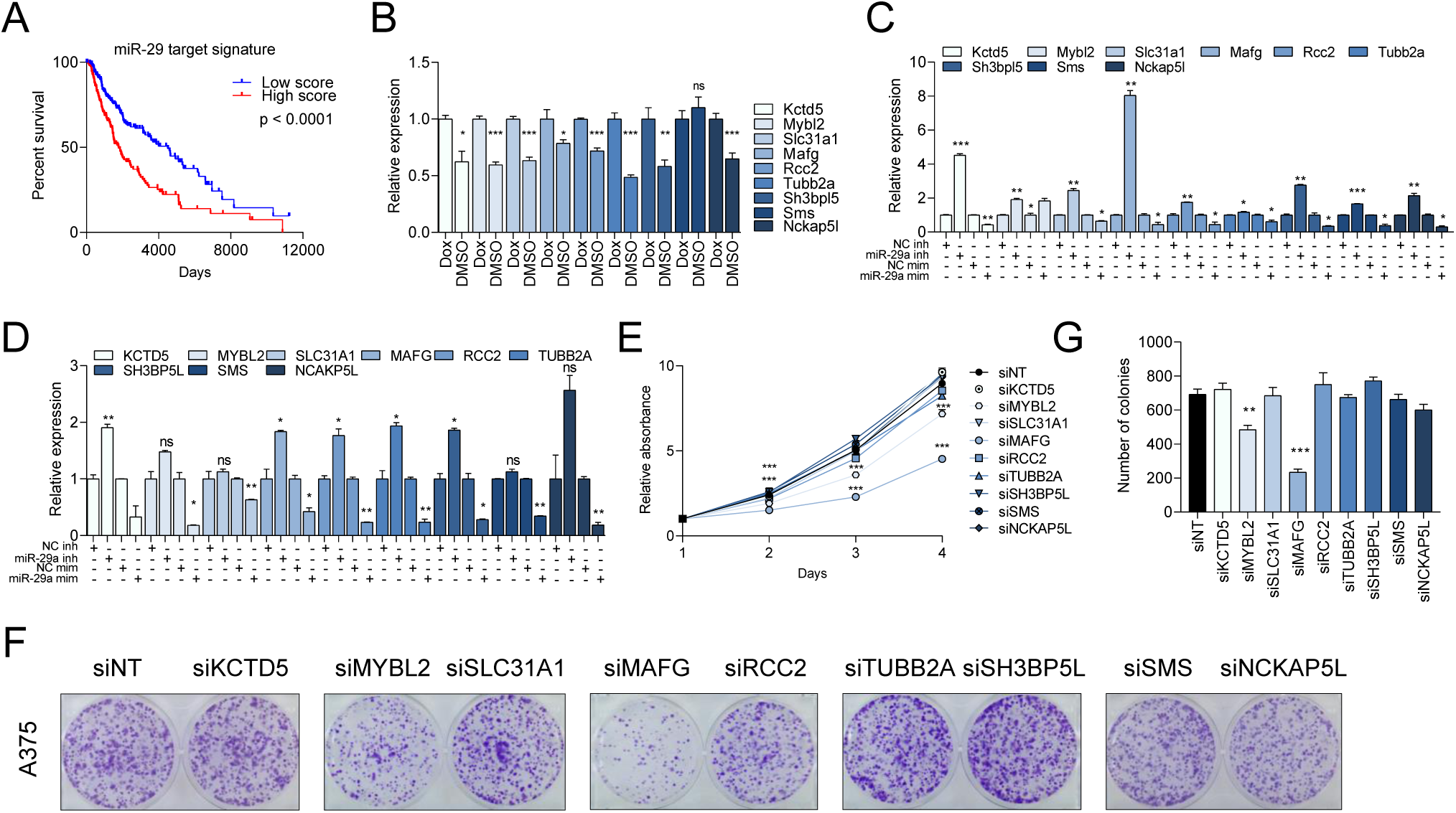
miR-29 targets constitute a vulnerability of melanoma. (A) Survival analysis from TCGA (PanCancer Atlas, n = 363) comparing melanoma patients with high or low expression of the 9 putative miR-29 targets. (B-D) Expression of the nine putative miR-29 target genes upon miR-29 reactivation in miR-29 sponge melanoma cells (B), and miR-29 inhibitor or mimic transfection in miR-29 sponge melanoma cells (C) or human A375 melanoma cells (D). The mean ± SEM of one representative out of three independent experiments performed in triplicate is shown. Gene expression levels are normalized to β-Actin (mouse) or GAPDH (human) as endogenous control. (E-G) Effect of the silencing of the nine putative miR-29 targets in the human melanoma A375 cell line on proliferation (E) and focus formation (F, G). The mean ± SEM of one representative out of two independent experiments performed in quadruplicate is shown, ns, not significant; * p < 0.05; ** p < 0.01; *** p < 0.001.

### MAFG is a bona fide target of miR-29 in melanocytes and melanoma

To validate MAFG as a target of miR-29 we transfected miR-29a, miR-29b, or miR-29c mimics into BRAF^V600E^-expressing melanocytes (H1B) and melanoma cells (WM164). We observed a general reduction of MAFG mRNA and protein expression (Figure 5A and 5B). By contrast, hairpin inhibitors of miR-29a, miR-29b, or miR-29c increased MAFG mRNA and protein levels in these cell lines (Figure 5A and 5B). Next, we created a MAFG 3’UTR Luciferase reporter and mutated the seed sequence of the miR-29 binding site with the highest prediction score (Supplementary Figure 5A). Co-transfection of miR-29 inhibitor with the wildtype MAFG 3’UTR reporter into H1B, H3B8, A375, and WM164 cells increased Luciferase activity (Figure 5C and Supplementary Figure 5B). By contrast, miR-29 mimics reduced the activity of the wildtype MAFG 3’UTR reporter and this effect was partially rescued by the miR-29 binding site mutation (Figure 5D and Supplementary Figure 5C). These findings indicate that MAFG is a *bona fide* target of miR-29.

**Figure 5:**
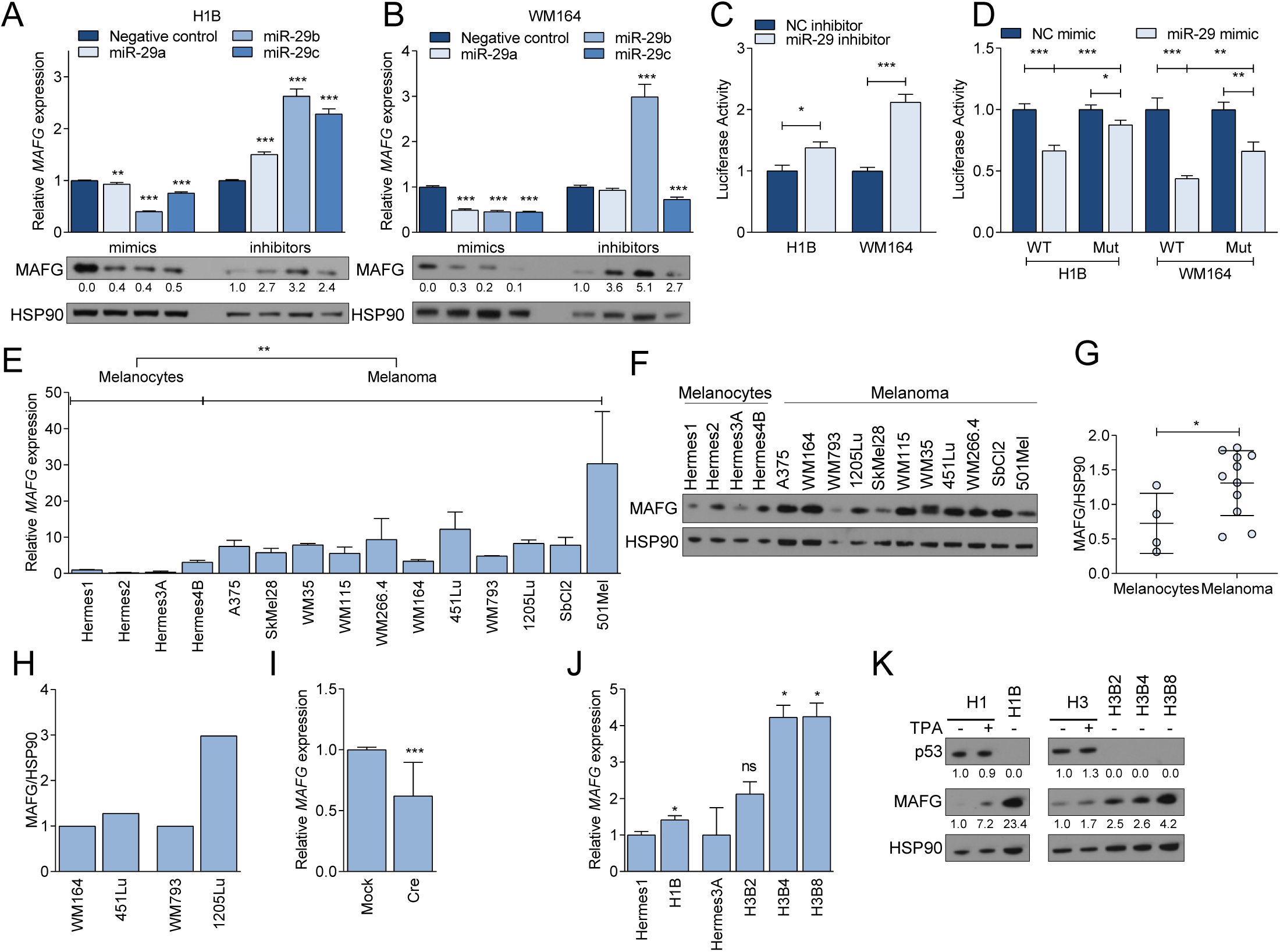
MAFG is directly regulated by miR-29. (A and B) MAFG mRNA (upper panels) and protein (lower panels) expression upon miR-29 mimics (left) or inhibitors (right) transfection in H1B (A) or WM164 (B). The mean ± SEM of one representative out of two independent experiments performed in triplicate is shown. Gene expression levels are normalized to GAPDH as endogenous control. (C) Activity of MAFG 3’UTR Luciferase reporter in response to miR-29 inhibitors. (D) Activity of MAFG wildtype or miR-29 binding site-mutant 3’UTR Luciferase reporter in response to miR-29 mimics. For (C) and (D) the combined mean ± SEM of two independent experiments performed in quadruplicate is shown. (E) qRT-PCR showing the basal expression levels of MAFG mRNA in melanocytes and melanoma cells. The mean ± SEM of one representative out of two independent experiments performed in triplicate is shown. Gene expression levels are normalized to GAPDH as endogenous control. (F) Western blot showing the basal expression levels of MAFG protein in melanocytes (n = 4) and melanoma cells (n = 11). (G) Quantification of the Western blot shown in (F). (H) Protein levels of MAFG in primary vs. metastatic cells relative to HSP90. (I) qRT-PCRs showing the expression of MAFG in LSL-Braf^V600E^ PMM following Adeno-Cre/Mock infection. **(J)** qRT-PCR showing the basal expression levels of MAFG mRNA in parental and BRAF^V600E^-mutant melanocytes. The mean ± SEM of one representative out of two independent experiments performed in triplicate is shown. Gene expression levels are normalized to GAPDH as endogenous control. (K) Western blot showing MAFG expression in response to TPA stimulation or BRAF^V600E^ expression. The mean ± SEM of one representative out of two independent experiments performed in triplicate is shown. Gene expression levels are normalized to GAPDH as endogenous control. * p < 0.05; ** p < 0.01; *** p < 0.001.

We next assessed if the expression of MAFG is altered during melanoma development First, we analyzed MAFG expression in melanocytes and melanoma cell lines and found that MAFG mRNA levels are increased in melanoma cell lines (Figure 5E). This finding corroborates the increase in MAFG mRNA observed in primary melanomas compared to nevi (Supplementary Figure 4D). MAFG protein expression was similarly elevated in melanoma cell lines compared to melanocytes (Figure 5F and 5G) and the metastatic derivatives 451Lu and 1205Lu showed increased MAFG protein levels compared to their parental cell lines WM164 and WM793, respectively (Figure 5H). Notably, acute activation of endogenous Braf^V600E^ in primary melanocytes diminished *MAFG* mRNA expression (Figure 5l), which correlated with increased miR-29 expression (Figure 2A). Conversely, *MAFG* mRNA levels were increased in BRAF^V600E^-expressing Hermes cells in which p53 is lost and pri-miR-29b2∼c is reduced (Figure 5J). This increase in *MAFG* mRNA led to a robust increase in MAFG protein expression that is more pronounced than stabilizing MAFG protein through TPA-induced MAPK signaling (Figure 5K).

### MAFG is critical for melanoma cell proliferation and has oncogenic potential

We next examined the role of MAFG in regulating melanocyte transformation and melanoma cell growth. In addition to A375 cells, silencing of MAFG decreased proliferation and focus formation in WM164, WM793, and WM266.4 melanoma cell lines (Supplementary Figures 6A-6F). Silencing of MAFG reduced proliferation of Hermes1 and Hermes3A melanocyte cell lines, but had no effect on Hermes2 and Hermes4B (Supplementary Figure 6G-6K). Moreover, silencing of the closely related family members MAFF and MAFK had no effect on proliferation or focus formation of A375 and WM164 cells (Supplementary Figure 7A-7E). Thus, MAFG is critical for melanoma cell growth.

To further test its oncogenic potential, we overexpressed MAFG in melanocytes. Constitutive overexpression of lentiviral MAFG in Hermes1 cells increased proliferation (Figure 6A) and conferred the ability to form colonies (Figure 6B). MAFG acts as a heterodimeric transcription factor; however, the high MAFG levels achieved by constitutive overexpression could promote homodimer formation with dominant negative effects. Therefore, we also used a Dox-inducible MAFG overexpression approach, which enabled the titration of MAFG levels. Moderate MAFG overexpression in Hermes1 (Figure 6C) increased proliferation and focus formation under regular (10% FBS) and low (2.5% FBS) serum conditions (Figure 6D–6F). Similarly, MAFG overexpression in WM164 cells modestly increased proliferation and augmented the focus formation capacity (Supplementary Figure 7F-I). To examine whether miR-29 elicits its tumor suppressive effects by repressing MAFG, we conducted a rescue experiment To this end, we transfected miR-29 mimics into MAFG-overexpressing or control Hermes1 cells. As expected, miR-29 mimics reduced proliferation and low-density clonogenic growth (Figure 6G and 6H). Importantly, the effects of miR-29 on proliferation and low-density growth were completely rescued by MAFG overexpression, indicating that miR-29 suppresses melanoma, at least in part, by repressing MAFG.

**Figure 6:**
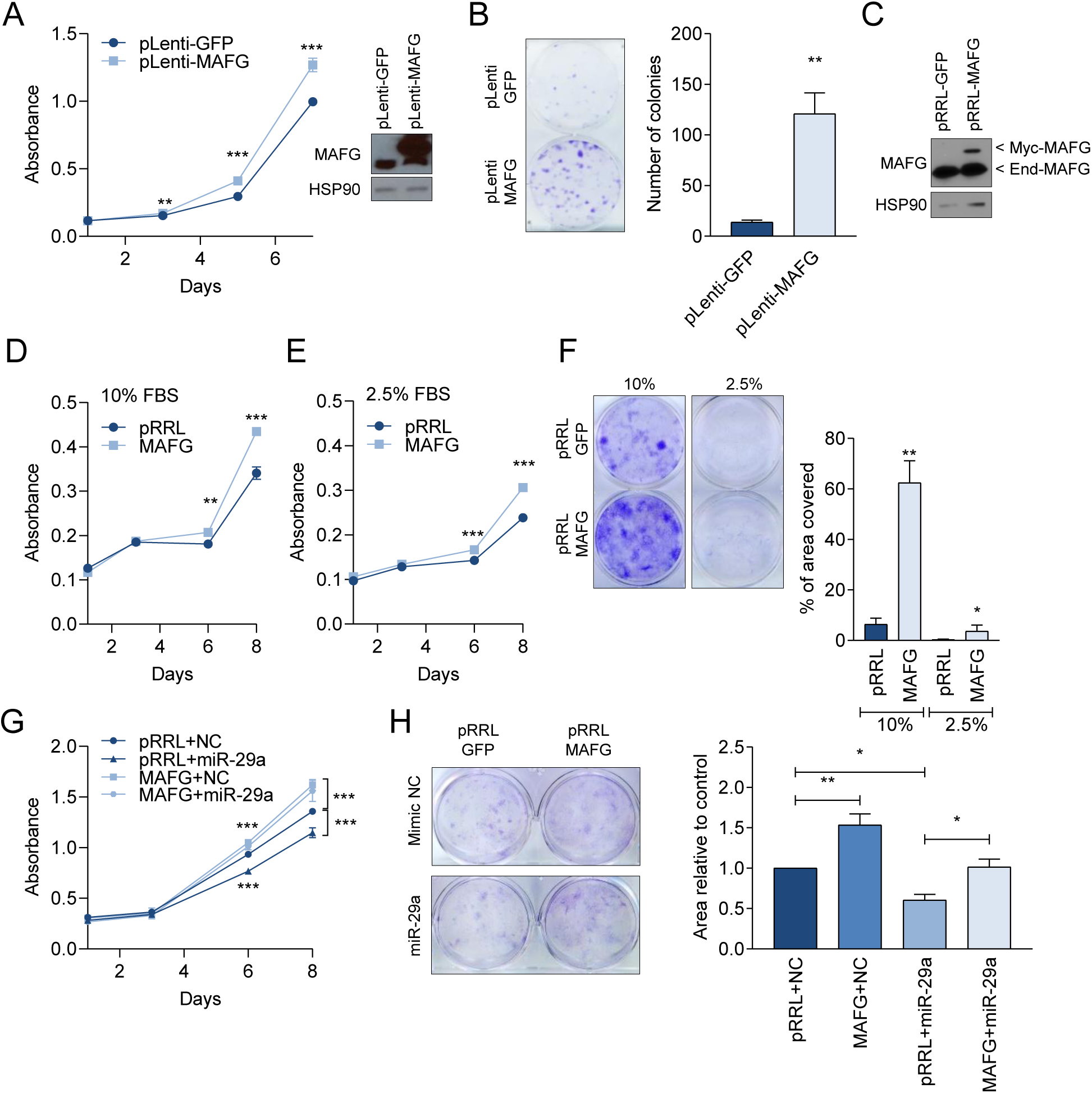
MAFG is a putative melanoma oncogene. (A, B) The effect of constitutive MAFG overexpression in Hermes1 cells on proliferation (A) and colony formation (B). (C) Validation of MAFG overexpression by a Dox-inducible vector in Hermes1 cells. (D-F) Proliferation (D, E) and colony formation (F) of Hermes1 in 10% FBS (D, F) or 2.5% FBS (E, F) cells upon acute and moderate MAFG induction by Dox. (G-H) Proliferation (G) and focus formation (F) of Hermes1 overexpressing MAFG in combination with miR-29 mimics. The mean ± SEM of two independent experiments performed in quadruplicate is shown. Western blots in (A and C) show the MAFG protein expression changes. For all Western blots, HSP90 was used as loading control. * p < 0.05; ** p < 0.01; ***p< 0.001.

## DISCUSSION

Deregulation of miRNAs frequently occurs in cancer and is thought to play critical roles in all aspects of tumorigenesis. Here, we investigated the deregulation of miR-29 in melanoma formation. Using MEFs and human melanocytes, we uncovered paradoxical upregulation of miR-29b1∼a by oncogenic RAS and BRAF, while MAPK signaling and p53 act in concert to promote miR-29b2∼c expression. Diminished expression of the p53-dependent miR-29b2∼c cluster is associated with the progression of nevi to frank melanoma, and inactivation of miR-29 promotes melanoma development in mice. De-repression of MAFG, which we identified as a *bona fide* target of miR-29, may contribute to melanoma development

Previous studies have shown that p53 regulates the expression of both miR-29b1∼a and miR-29b2∼c (Ugalde et al. 2011; Chen et al. 2018). However, our results indicate that transcription of miR-29b1∼a is independent of p53, both in MEFs and in melanocytes. Rather, miR-29b1∼a is regulated directly via MAPK signaling. This discrepancy is likely due to the fact that mature miR-29 species were analyzed by qRT-PCR in the previous studies, a method that failed to distinguish mature miR-29 family members, as has been suggested previously (Kurinna et al. 2014). We also observed regulation of miR-29b2∼c by MAPK signaling; however, this requires expression of p53. It remains to be investigated how MAPK signaling and p53 activation coordinately enhance miR-29b2∼c expression upon acquisition of an oncogenic BRAF mutation. p53 may be induced by oncogene-activated MAPK pathway hyperactivation as ERK has been reported to phosphorylate p53 at serine 15 (Wu 2004). Moreover, MAPK activation increases expression of Cyclin D, which releases E2F-1 via RB phosphorylation, thereby promoting pl4ARF transcription (Peeper et al. 1997; Bates et al. 1998). pl4ARF in turn stabilizes p53 through inhibition of MDM2 (Sherr and Weber 2000). MAPK hyperactivation downstream of oncogenic BRAF is a critical driver of melanoma development (Davies et al. 2002), and p53 activation in response to mutant BRAF has been observed in melanocytes (Yu et al. 2009; Ko et al. 2019). Thus, oncogenic BRAF may activate p53 in melanocytes via the MAPK pathway, which in turn induces pri-miR-29b2∼c expression. Alternatively, MAPK signaling may directly promote pri-miR-29b2∼c expression, and p53 serves as a critical permissive factor that is required for pri-miR-29b2∼c transcription. Oncogenic BRAF only very moderately activates p53 (Lloyd et al. 1997; Sheu et al. 2012), suggesting that BRAF may indeed work in concert with rather than through p53.

Several reports describe tumor suppressive functions of miR-29 in cultured cells, including in melanoma cell lines (Nguyen et al. 2011; Yu et al. 2014; Cui et al. 2015; Nishikawa et al. 2015; Chen et al. 2018; Xiong et al. 2018), which we corroborated in our study. Given the tumor suppressive functions of miR-29 and its regulation by MAPK signaling, we hypothesized that MAPK hyperactivation could provoke a miR-29-dependent barrier that prevents melanoma formation. The MAPK pathway is almost universally hyperactivated in melanoma, owing to the frequent activating mutations in BRAF and NRAS (Davies et al. 2002; Satyamoorthy et al. 2003; Gray-Schopfer et al. 2005; Sumimoto et al. 2006). Notably, growth arrested nevi are common in humans and >80% of nevi harbor BRAF^V600E^ mutations (Wu et al. 2007; Yeh et al. 2013), suggesting that such a barrier indeed exists (Michaloglou et al. 2005). To overcome this barrier, BRAF/NRAS mutant melanocytes must reverse the increase in miR-29 levels. We observed that miR-29b2∼c expression is decreased i) in melanocytes upon chronic expression of BRAF^V600E^, which also provoked loss of p53, ii) in melanoma cell lines compared to melanocytes, and iii) in primary melanomas compared to nevi. It is tempting to speculate that while miR-29b1∼a remains elevated due to continuous MAPK hyperactivation, impaired p53 activity leads to decreased miR-29b2∼c expression, thereby promoting progression from nevi to frank melanoma (Figure 7). pl6INK4A was initially believed to be the main mediator of oncogene-induced senescence in melanocytes. However, deletion of pl6Ink4a in a BRAF^V600E^ melanoma mouse model fails to provoke broad circumvention of melanocyte senescence (Dhomen et al. 2009), and a more complex regulation of melanocyte senescence involving p53 has emerged (Bansal and Nikiforov 2010). Indeed, p53 may play a role in the growth arrest of nevi (Gray-Schopfer et al. 2006; Yu et al. 2009; Terzian et al. 2010), and p53 inactivation in genetically engineered mice promotes melanoma development in the context of BRAF^V600E^ (Viros et al. 2014). Moreover, p53 mutations and copy number losses occur frequently in cutaneous melanoma (Berger et al. 2012; Hodis et al. 2012; Krauthammer et al. 2012; Krauthammer et al. 2015; Zhang et al. 2016). Even more common are deletions of CDKN2A (Hodis et al. 2012; Krauthammer et al. 2012; Krauthammer et al. 2015; Zhang et al. 2016), which besides pl6INK4A encodes ARF, a positive regulator of p53 protein stability. A pl6INK4A-independent role for ARF in melanoma suppression has been described (Hewitt et al. 2002; Sharpless et al. 2003; Freedberg et al. 2008). Additionally, a subset of melanomas harbor amplifications of MDM2 (Muthusamy et al. 2006; Cancer Genome Atlas 2015), an E3 ubiquitin ligase that promotes the turnover of p53. Thus, multiple mechanisms of p53 inactivation occur in melanoma, all of which could lead to a reduction in miR-29b2∼c expression.

**Figure 7:**
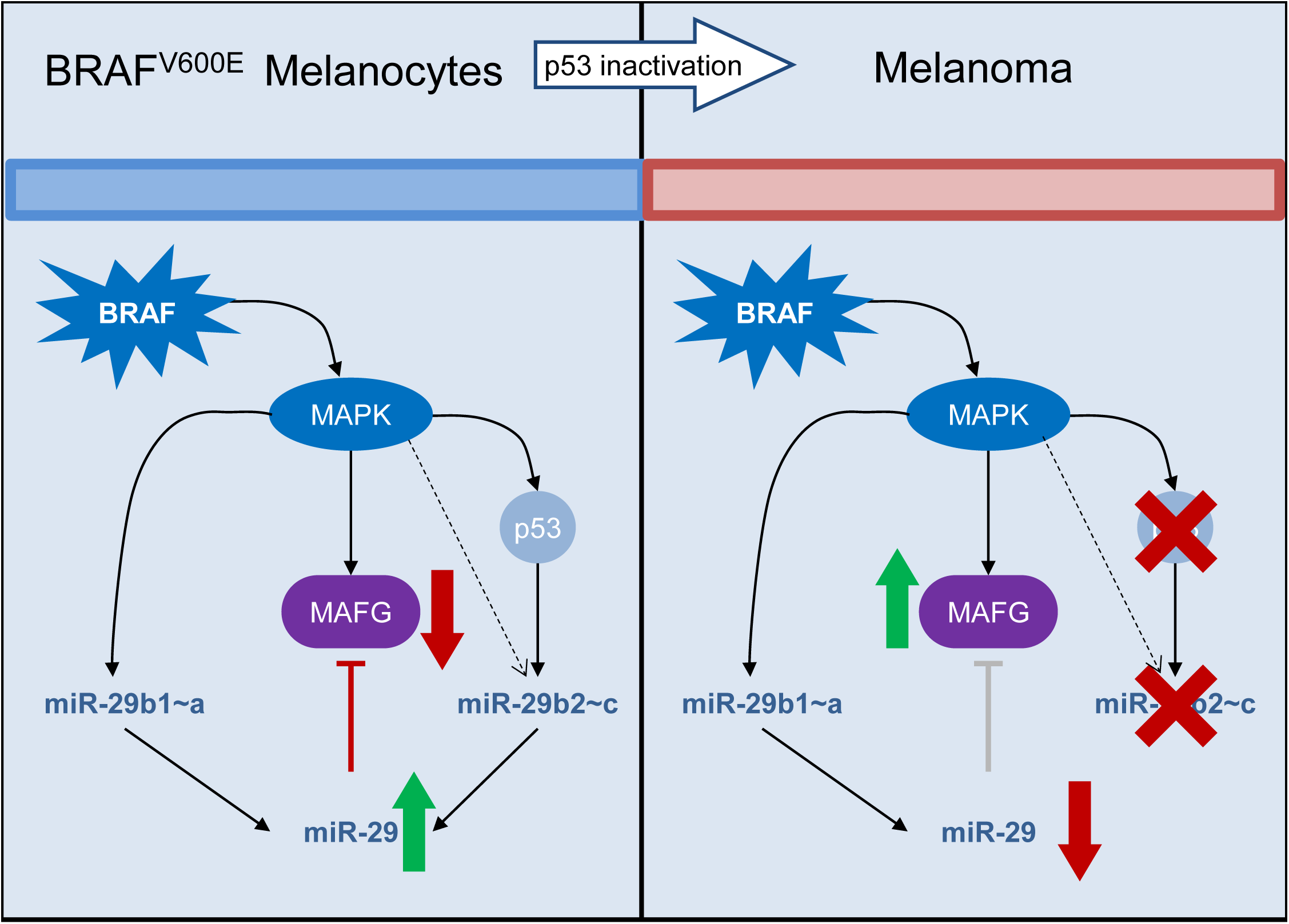
MAPK-induced miR-29 targets MAFG and suppresses melanoma development. Oncogenic MAPK signaling in nevi stimulates transcription of pri-miR-29b1∼a and pri-miR-29b2∼c in a p53-independent and p53-dependent manner, respectively, resulting in MAFG repression. Inactivation of p53 during the progression from nevi to frank melanoma decreases the p53-dependent transcription of pri-miR-29b2∼c, thereby leading to de-repression of MAFG and augmented melanoma development.

Using a high-throughput mouse modeling approach, we inactivated miR-29 through expression of a sponge construct specifically in Braf^V600E^; Pten^Δ/WT-^ melanocytes. In agreement with our hypothesis, miR-29 inactivation accelerated the development of melanoma. Not only is this the first model used to study miR-29 inactivation in tumorigenesis, our approach also affirms that synthetic miRNA sponges are powerful tools to examine miRNA function *in vivo*. One advantage over traditional modeling approaches, such as the previously published conditional knock-out allele of miR-29b1∼a (Kogure et al. 2012), is that a miRNA sponge has the potential to inactivate all members of a miRNA family. However, it is usually not clear how well a sponge interacts with each family member, especially in cases like the miR-29 family where one member, miR-29b, also localizes to the nucleus (Hwang et al. 2007). Thus, future studies using alternative approaches such as CRISPR/Cas9-mediated specific deletion of individual clusters or miRNAs will further elucidate the role of each miR-29 family member in melanoma. This will also address if a decrease in miR-29b2∼c in melanoma simply lowers the overall miR-29 levels to promote melanoma, indicative of functional redundancy of miR-29a, −29b, and −29c, or if miR-29b and miR-29c have specific functions that are impaired upon miR-29b2∼c downregulation.

Since we did not observe changes in the expression of the validated miR-29 targets *AKT3, MCL1*, and *DNMT3B*, we identified new targets whose repression may contribute to restricting the progression of nevi to melanoma. Of the nine genes identified by our approach, RCC2, MYBL2 and SLC31A1 had previously been identified as miR-29 targets (Martinez et al. 2011; Matsuo et al. 2013; Wu et al. 2013; Hu et al. 2014; Sun et al. 2018). While all nine genes were validated as miR-29 targets and the miR-29 sponge modulated the expression of eight of these targets, silencing of only MAFG and MYBL2 diminished the proliferation of melanoma cells. Thus, miR-29-mediated repression of

MAFG and/or MYBL2 may impair the transition from nevi to frank melanoma. We selected MAFG for further analyses because its silencing had a more severe effect on melanoma cells than MYBL2 knockdown. Interestingly, the MAFG protein is stabilized by ERK-mediated phosphorylation (Fang et al. 2016), suggesting that MAPK signaling converges on MAFG via ERK and miR-29. In addition to being repressed by miR-29, TCGA data indicate copy number gains of MAFG in melanoma. Thus, MAFG is deregulated in melanoma through multiple mechanisms, and our *in vitro* data further suggest that MAFG plays an important role in melanoma biology. Oncogenic roles for MAFG have so far been described in lung, ovarian, colorectal, and liver cancer (Fang et al. 2016; Vera et al. 2017; Liu et al. 2018), and, interestingly, MAFG is a binding partner of CNC and BACH protein families, including NRF2 (reviewed in (Katsuoka and Yamamoto 2016)). NRF2 is a critical regulator of redox biology and cellular metabolism (Hirotsu et al. 2012; Hayes and Dinkova-Kostova 2014), and mutations in NRF2 and its negative regulator KEAP1 occur frequently in lung and upper airway cancers (Cloer et al. 2019). By contrast, mutations in NRF2/KEAP1 are rarely observed in melanoma, but a recent study nevertheless suggests an important role for NRF2 in melanoma differentiation and interaction with the microenvironment (Jessen et al. 2020). Future studies will address if MAFG functions in melanoma through binding to and enhancing the transcriptional activity of NRF2, or if other binding partners are critical. Thus, our work has uncovered that miR-29 prevents MAPK pathway-driven melanoma formation, at least in part, by repressing MAFG.

## MATERIALS AND METHODS

### Cell culture and treatments

The human immortalized melanocytes cell lines Hermes1, Hermes2, Hermes3A, and Hermes4B were obtained from the Functional Genomics Cell Bank at St George’s, University of London, UK, and cultured in RPMI media supplemented with 10% FBS, 10ng/mL hSCF (R&D, Cat # 255-SC), 200nM TPA (Sigma, Cat # P8139), 200pM Cholera Toxin (Sigma, Cat # C8052), and 10nM Endothelin-1 (Sigma, Cat # E7764) at 37°C in a humidified atmosphere containing 10% CO_2_. A375 and SK-Mel28 cells were purchased from ATCC, WM164, WM35, WM793 and 1205Lu, WM115, WM266.4, 451Lu cells were a gift from M. Herlyn from the Wistar Collection of Melanoma cell lines, SbC12 were provided by D. Tuveson, and 501Mel were from K. Smalley. The colon cancer cell line HT29 was a gift from L. Dow and the H1755 and H1395 lung cancer cell lines were provided by G. DeNicola. All cancer cell lines were cultured in RPMI containing 5% FBS at 37°C in a humidified atmosphere containing 5% CO_2_. The BRAF^V600E^-expressing human melanocyte cell lines were derived by infecting Hermes1 and Hermes3A with a BRAF^V600E^-pLenti-Hygro (provided by L. Wan) in the presence of 8μg/mL Polybrene. Transduced cells were selected in 100μg/mL of Hygromycin (Invivogen, Cat # ant-hg-1) for seven days in the absence of TPA. Four independent clones were picked and expanded until stable cell lines were obtained. MEFs were generated from E13.5-E14.5 embryos from LSL-Braf^V600E^ (Karreth et al., Mol Cell 2009) or LSL-Kras^G12D^ mice (Jackson et al., Genes & Dev, 2001) and cultured in DMEM containing 10% FBS at 37°C in a humidified atmosphere containing 5% CO_2_. Wildtype MEFs were derived from littermate embryos that did not harbor the LSL-Braf^V600E^ or LSL-Kras^G12D^ alleles. LSL-Braf^V600E^ primary mouse melanocytes (PMM) were isolated as previously described (Yuspa and Harris 1974) with minor modifications. To recombine floxed alleles, MEFs and PMM were infected with approximately 10^7^pfu/mL Ad5CMVCre or Ad5CMVempty adenovirus obtained from the University of Iowa Viral Vector Core. Lenti-X 293T cells were obtained from Takara and cultured in DMEM containing 10% FBS at 37°C in a humidified atmosphere containing 5% CO_2_. All cell lines were routinely tested for mycoplasma using MycoAlert Plus (Lonza, Cat # LTO7-710), and human melanoma cell lines were STR authenticated by Moffitt’s Molecular Genomics Core. Doxorubicin (Fisher Scientific, Cat # BP25131) was used at a final concentration of ΙΟμΜ for 24 hours and AZD6244 (Selleckchem, Cat # S1008) was used at a final concentration of 0.5 μΜ for 8 or 24 hours.

### RNA isolation and quantitative RT-PCR

Total RNA was isolated using TRI-Reagent (Zymo Research, Cat # R2050-1-200) and mature miRNAs were isolated using the miRNeasy Mini Kit (Qiagen, Cat # 217004) according to the manufacturers’ recommendations. For qRT-PCR, 500ng of total RNA were retrotranscribed using PrimeScript RT Master Mix (Takara Bio, Cat. # RR036A), and subsequent TaqMan assay-based or SYBR Green-based qPCRs were performed using PerfeCTa qPCR ToughMix (QuantaBio, Cat. # 97065-960) or PerfeCTa SYBR Green FastMix (QuantaBio, Cat. # 95073-012), respectively. Mature miRNAs were retrotranscribed using TaqMan MicroRNA Reverse Transcription Kit (Thermo Fisher, Cat # 4366596) and analyzed by qPCR using PerfeCTa qPCR ToughMix (QuantaBio, Cat # 97065-960). Samples were analyzed in triplicate using the StepOne Plus PCR system (Applied Biosystems, USA). The comparative threshold cycle method (2^ΔΔCt^) was used to calculate the relative expression levels. snoU6 was used as endogenous control for mature miRNAs while GAPDH or β-Actin were used for mRNAs and pri-miRNAs.

TaqMan Probes for expression analyses were purchased from Thermo Fisher Scientific (U6 snRNA: 001973; mouse β-actin: Mm02619580_gl; human β-ACTIN: mouse Cdkn1a: Mm04205640_gl; mmu-mir-29a: Mm03306859_pri; mmu-mir-29b-2: Mm03307196_pri; mmu-mir-29c: Mm03306860; mmu-mir-29b-l; Mm03306189_pri; hsa-mir-29a: Hs03302672_pri; hsa-mir-29c: Hs04225365_pri; hsa-miR-29a: 002112; hsa-miR-29b: 000413; hsa-miR-29c: 000587). Primers for SYBR Green qPCR are listed in Supplementary Table 1.

### RNA-sequencing

Human melanoma cell lines and Hermes melanocyte cell lines were grown in RPMI medium containing 10% FBS in the absence of growth factors for 48 hours to ensure similar culture conditions for melanoma cells and melanocytes. Total RNA from cells was isolated using miRNeasy Mini Kit (Qiagen, Cat # 217004) and RIN were analyzed on an Agilent TapeStation.

#### Sequencing by the Biostatistics and Bioinformatics Core at Moffitt

Double-stranded cDNA synthesis and library construction were performed using the Nugen Ovation Human RNA-seq library system (Nugen, San Carlos, CA). An Illumina NextSeq 550 instrument was used to generate 75 base pair paired-end RNA-seq reads. The raw reads were first assessed for quality using FastQC (http://www.bioinformatics.babraham.ac.uk/projects/fastqc/) and then trimmed by cutadapt (Martin 2011) (version 1.8.1) to remove reads with adaptor contaminants and low-quality bases. Read pairs with either end too short (<25bps) were discarded from further analysis. Trimmed and filtered reads were aligned to the GRCh37 human reference genome using STAR (Dobin et al. 2013) (version 2.5.3a). Uniquely mapped reads were counted by htseq-count (Anders et al. 2015) (version 0.6.1) using Gencode v25 primary annotation. Differential expression analysis was performed using DESeq2 taking into account of RNA composition bias (Love et al. 2014). Genes with fold-changed and false discovery rate (FDR) q-values< 0.05 were considered differentially expressed.

#### Sequencing by Novogene (Sacramento, CA)

Library preparation was performed by NEBNext® Ultra^TM^ RNA Library Prep Kit for Illumina® which is for non-stranded libraries. Sequencing was performed on the NovaSeq 6000 with a paired end 150 BP method. Downstream analysis was performed using a combination of programs including STAR, HTseq, Cufflink and our wrapped scripts. Alignments were parsed using Tophat program and differential expressions were determined through DESeq2/edgeR. GO and KEGG enrichment were implemented by the ClusterProfiler. Gene fusion and difference of alternative splicing event were detected by Star-fusion and rMATS software. Reference genome and gene model annotation files were downloaded from genome website browser (NCBI/UCSC/Ensembl) directly. Indexes of the reference genome was built using STAR and paired-end clean reads were aligned to the reference genome using STAR (v2.5). STAR used the method of Maximal Mappable Prefix(MMP) which can generate a precise mapping result for junction reads. HTSeq vO.6.1 was used to count the read numbers mapped of each gene. And then FPKM of each gene was calculated based on the length of the gene and reads count mapped to this gene. FPKM, Reads Per Kilobase of exon model per Million mapped reads, considers the effect of sequencing depth and gene length for the reads count at the same time, and is currently the most commonly used method for estimating gene expression levels (Mortazavi et al. 2008). Differential expression analysis between was performed using the DESeq2 R package (2_1.6.3). DESeq2 provide statistical routines for determining differential expression in digital gene expression data using a model based on the negative binomial distribution. The resulting P-values were adjusted using the Benjamini and Hochberg’s approach for controlling the False Discovery Rate (FDR). Genes with an adjusted P-value < 0.05 found by DESeq2 were assigned as differentially expressed. Accession numbers: GSE148552, PRJNA624657

#### Plasmids

pBabe, pBabe-Braf^V600E^, and pBabe-Kras^G12D^ were gifts from D. Tuveson. pLenti-GFP-puro was purchased from Addgene (plasmid #17448). The CMV promoter and puromycin in pLenti-GFP-puro were replaced with the EF1α promoter and blasticidin, respectively, using standard In-Fusion cloning (Takara Bio, Cat # 638911) to create pLEGB. The Myc-DDK-tagged ORF clone of MAFG (RC221486, OriGene USA) was a gift from I. Ibanez de Caceres and cloned into pLEGB to replace GFP. The full length MAFG 3’UTR sequence (NM_002359.3 OriGene, USA) was a gift from I. Ibanez de Caceres and was cloned into psiCHECK2 plasmid by In-Fusion cloning. The psiCHECK2-miR29 Luciferase reporter was created by oligo cloning into psiCHECK2. For inducible expression, MAFG was cloned into the Dox-inducible pRRL-Blast vector by In-Fusion cloning. All primers for cloning are detailed in Supplementary Table 1.

### Site-directed mutagenesis

The psiCHECK2-MAFG-3’-UTR reporter plasmid was used to generate the miR-29 binding site mutant. Four different miR-29 binding sites were predicted by TargetScan, one of which is highly conserved. Site-directed mutagenesis was performed using the Q5 Site-Directed Mutagenesis Kit according to the manufacturer’s instructions. Primers for site-directed mutagenesis are listed in Supplementary Table 1

### Cell transfection and lentiviral transduction

For miR-29 overexpression and inhibition, 100,000 cells/well were plated in 6-well plates and transfected with 25 to 150 nM of Dharmacon miRIDIAN microRNA miR-29a, miR-29b or miR-29c mimic (C-310521-07-0002, C-310381-05-0002 or C-310522-05-0002), hairpin inhibitor (IH-310521-08-0002, IH-310381-07-0002 or IH-310522-08-0002), or negative controls (CN-002000-01-05; IN-001005-01-05) using JetPrime (VWR Cat # 89129-924) according to the manufacturer’s protocol and assayed after 48 hours. For Luciferase assays, cells were plated in 96-well plates at a density of 10,000 cells/well. psiCHECK2-MAFG-3’UTR_wildtype or psiCHECK2-MAFG-3’UTR_miR-29-mutant were co-transfected with miR-29 mimics or inhibitors, following the procedure described above. Luminescence was assayed after 24 hours using the Dual Luciferase Assay System (Promega, Cat # E1960), according to the manufacturer’s instructions. Similarly, miR-29-sponge melanoma cell lines were plated at a density of 10,000 cells/well and transfected with psiCHECK2 or psiCHECK2-miR-29 reporter following the same procedure. Luminescence was assayed after 48 hours using the Dual Luciferase Assay System. Results were normalized to Firefly luminescence. For retroviral transductions, Lenti-X 293T cells were transfected with the retroviral vector and Eco helper plasmid at a 2:1 ratio. For lentiviral transductions, Lenti-X 293T cells were transfected with the lentiviral vector and the Δ8.2 and pMD2-VSV-G helper plasmids at a 9:8:1 ratio. Supernatants were collected 48 hours after transfection and filtered through a 0.45μm filter. Cells were plated in 6-well plates at a density of 300,000 cells/well and transduced with supernatants in the presence of 8 μg/mL polybrene for 6 hours. Selection was carried out by treating cells with 10 μg/mL Blasticidin for 5 days or 1 μg/mL Puromycin for 4 days. For siRNA transfections, 100.0 cells/well were plated in 6-well plates and transfected with 25nM of ON-TARGETplus siRNA pool or Non-Targeting control using JetPrime according to the manufacturer’s protocol. 4-6 hours after transfection, cells were trypsinized and replated for cell biological assays. siRNA pool catalog numbers are available in Supplementary Table 1.

### Proliferation and colony formation assays

For proliferation assays, cells were plated in 96-well plates at a density of 1,000–2,500 cells/well and harvested for five days. Cells were fixed and stained with 0.1% crystal violet (VWR, Cat # 97061-850) solution in 20% methanol for 20 minutes followed by extraction of crystal violet with 10% acetic acid. Absorbance was measured at 600 nm using a plate reader. For colony formation assays, cells were plated in 6-well plates at a density of 1,000–2,000 cells/well and cultured for 2-3 weeks. Cells were fixed and stained with 0.1% crystal violet (VWR, Cat# 97061-850) solution in 20% methanol for 20 minutes. Colonies were quantified using Image J software.

### Immunoblotting

Protein isolation was performed as previously described (Bok et al. 2019). 20μg of total protein were subjected to SDS-PAGE and Western blot, performed as previously described (Bok et al. 2019). Primary antibodies used were BRAF (Sigma, Cat # HPA001328), KRAS (Santa Cruz, Cat # sc-30), HA-Tag (Cell Signaling, Cat # 3724T), human p53 (Santa Cruz, Cat # sc-126), mouse p53 (BioVision, Cat # 3036-100), p21 (Abeam, Cat # ab109199), ERK (Cell Signaling, Cat # 4695), pERK (Cell Signaling Cat # 9101S), c-Jun (Cell Signaling Cat # 9165S), MAFG (Thermo Fisher Scientific, Cat # PAS-90907) and HSP90 (Cell Signaling, Cat # 4874)

### ES cell targeting, mouse generation, and ESC-GEMM experiments

ES cell targeting and generation of chimeras was performed a described previously (Bok et al. 2019). Melanoma development was induced in 3-4 week old miR-29 sponge and GFP control chimeras having similar ESC contribution using 25mg/mL 4-OH Tamoxifen as described previously (Bok et al. 2019). Mice were fed 200mg/kg Doxycycline (Envigo, Cat # TD180625) *ad libitum*. All animal experiments were conducted in accordance with an IACUC protocol approved by the University of South Florida. Experimental mice were euthanized when IACUC-approved clinical endpoints, typically volume of primary tumors, was reached. The derivation of the murine melanoma cell line from an ESC-GEMM chimera was performed as described previously (Bok et al. 2019).

### Analysis of miR-29 target expression in The Cancer Genome Atlas

We obtained mRNA expression and survival data for KCTD5, MYBL2, SLC31A1, MAFG, RCC2, TUBB2A, SH3BP5L, SMS, and NCKAP5L of 354 skin cancer melanoma tumors from The Cancer Genome Atlas (SKCM-TCGA). To generate a scoring for all nine genes with equal contribution, we normalized the mRNA expression values to the average for each gene, followed by calculating the average of each gene for each patient For survival analysis, the data was stratified for patients with high or low score according to the median (cutoff: 0.96) and overall survival was estimated according to the Kaplan-Meier method. Groups were compared by the Log Rank test.

### Statistical analysis

Statistical analysis was performed using GraphPad Prism software. Survival data were compared by applying the Gehan-Breslow-Wilcoxon test, and all other data were analyzed with the unpaired two-tailed t-test or ordinary one-way ANOVA. A p-value below 0.05 was considered statistically significant. Data represent the mean ± SEM of at least two independent experiments performed at least in triplicate.

## ACKNOWLEDGEMENTS

We thank A. Nyein and O. Balogun for technical assistance and Dr. I. Ibanez-de-Caceres for plasmids. This work was supported by grants to F.A.K. from the NIH/NCI (R03 CA227349) and the Melanoma Research Alliance (MRA Young Investigator Award and Team Science Award). This work was also supported by the Gene Targeting Core, Molecular Genomics Core, and Biostatistics and Bioinformatics Core, which are funded in part by Moffitt’s Cancer Center Support Grant (NCI, P30-CA076292).

## AUTHOR CONTRIBUTIONS

Conceptualization, FAK and OV; Methodology, FAK, OV, IB; Investigation, FAK, OV, IB, NJ, KN, XX, NM, AA, KW and KYT; Writing-Original Draft, FAK and OV; Writing -Reviewing and Editing, FAK, OV, IB, NJ, KN, XX, NM, AA, KW and KYT. Funding Acquisition, FAK; Resources, FAK; Supervision FAK.

## REFERENCES

Alizadeh M, Safarzadeh A, Beyranvand F, Ahmadpour F, Hajiasgharzadeh K, Baghbanzadeh A, Baradaran B. 2019. The potential role of miR-29 in health and cancer diagnosis, prognosis, and therapy. Journal of cellular physiology 234: 19280–19297.

Anders S, Pyl PT, Huber W. 2015. HTSeq-a Python framework to work with high-throughput sequencing data. Bioinformatics 31: 166–169.

Bae HJ, Noh JH, Kim JK, Eun JW, Jung KH, Kim MG, Chang YG, Shen Q, Kim SJ, Park WS et al. 2014. MicroRNA-29c functions as a tumor suppressor by direct targeting oncogenic SIRT1 in hepatocellular carcinoma. Oncogene 33: 2557–2567.

Bansal R, Nikiforov MA. 2010. Pathways of oncogene-induced senescence in human melanocytic cells. Cell Cycle 9: 2782–2788.

Bates S, Phillips AC, Clark PA, Stott F, Peters G, Ludwig RL, Vousden KH. 1998. pl4ARF links the tumour suppressors RB and p53. Nature 395: 124–125.

Berger MF, Hodis E, Heffernan TP, Deribe YL, Lawrence MS, Protopopov A, Ivanova E, Watson IR, Nickerson E, Ghosh P et al. 2012. Melanoma genome sequencing reveals frequent PREX2 mutations. Nature 485: 502–506.

Bok I, Vera O, Xu X, Jasani N, Nakamura K, Reff J, Nenci A, Gonzalez JG, Karreth FA. 2019. A versatile ES cell-based melanoma mouse modeling platform. Cancer Res.

Cancer Genome Atlas N. 2015. Genomic Classification of Cutaneous Melanoma. Cell 161: 1681–1696.

Cloer EW, Goldfarb D, Schrank TP, Weissman BE, Major MB. 2019. NRF2 Activation in Cancer: From DNAto Protein. Cancer Res 79: 889–898.

Cui H, Wang L, Gong P, Zhao C, Zhang S, Zhang K, Zhou R, Zhao Z, Fan H. 2015. Deregulation between miR-29b/c and DNMT3A is associated with epigenetic silencing of the CDH1 gene, affecting cell migration and invasion in gastric cancer. PLoSOne 10: e0123926.

Chang TC, Yu D, Lee YS, Wentzel EA, Arking DE, West KM, Dang CV, Thomas-Tikhonenko A, Mendell JT. 2008. Widespread microRNA repression by Myc contributes to tumorigenesis. Nature genetics 40: 43–50.

Chen B, Wang J, Wang J, Wang H, Gu X, Tang L, Feng X. 2018. A regulatory circuitry comprising TP53, miR-29 family, and SETDB1 in non-small cell lung cancer. Bioscience reports 38.

Davies H, Bignell GR, Cox C, Stephens P, Edkins S, Clegg S, Teague J, Woffendin H, Garnett MJ, Bottomley W et al. 2002. Mutations of the BRAF gene in human cancer. Nature 417: 949–954.

Davis BN, Hata A. 2009. Regulation of MicroRNA Biogenesis: A miRiad of mechanisms. Cell communication and signaling: CCS 7: 18.

Dhomen N, Reis-Filho JS, da Rocha Dias S, Hayward R, Savage K, Delmas V, Larue L, Pritchard C, Marais R. 2009. Oncogenic Braf induces melanocyte senescence and melanoma in mice. Cancer cell 15: 294–303.

Dobin A, Davis CA, Schlesinger F, Drenkow J, Zaleski C, Jha S, Batut P, Chaisson M, Gingeras TR. 2013. STAR: ultrafast universal RNA-seq aligner. Bioinformatics 29: 15–21.

Fang M, Hutchinson L, Deng A, Green MR. 2016. Common BRAF(V600E)-directed pathway mediates widespread epigenetic silencing in colorectal cancer and melanoma. Proc Natl Acad Sci USA 113: 1250–1255.

Fattore L, Costantini S, Malpicci D, Ruggiero CF, Ascierto PA, Croce CM, Mancini R, Ciliberto G. 2017. MicroRNAs in melanoma development and resistance to target therapy. Oncotarget 8: 22262–22278.

Freedberg DE, Rigas SH, Russak J, Gai W, Kaplow M, Osman I, Turner F, Randerson-Moor JA, Houghton A, Busam K et al. 2008. Frequent pl6-independent inactivation of pl4ARF in human melanoma. Journal of the National Cancer Institute 100: 784–795.

Gray-Schopfer VC, Cheong SC, Chong H, Chow J, Moss T, Abdel-Malek ZA, Marais R, Wynford-Thomas D, Bennett DC. 2006. Cellular senescence in naevi and immortalisation in melanoma: a role for pl6? Br J Cancer 95: 496–505.

Gray-Schopfer VC, da Rocha Dias S, Marais R. 2005. The role of B-RAF in melanoma. Cancer metastasis reviews 24: 165–183.

Havelange V, Stauffer N, Heaphy CC, Volinia S, Andreeff M, Marcucci G, Croce CM, Garzon R. 2011. Functional implications of microRNAs in acute myeloid leukemia by integrating microRNA and messenger RNA expression profiling. Cancer 117: 4696–4706.

Hayes JD, Dinkova-Kostova AT. 2014. The Nrf2 regulatory network provides an interface between redox and intermediary metabolism. Trends in biochemical sciences 39: 199–218.

Hewitt C, Lee Wu C, Evans G, Howell A, Elles RG, Jordan R, Sloan P, Read AP, Thakker N. 2002. Germline mutation of ARF in a melanoma kindred. Hum Mol Genet 11: 1273–1279.

Hirotsu Y, Katsuoka F, Funayama R, Nagashima T, Nishida Y, Nakayama K, Engel JD, Yamamoto M. 2012. Nrf2-MafG heterodimers contribute globally to antioxidant and metabolic networks. Nucleic Acids Res 40: 10228–10239.

Hodis E, Watson IR, Kryukov GV, Arold ST, Imielinski M, Theurillat JP, Nickerson E, Auclair D, Li L, Place C et al. 2012. A landscape of driver mutations in melanoma. Cell 150: 251–263.

Hu Z, Klein JD, Mitch WE, Zhang L, Martinez I, Wang XH. 2014. MicroRNA-29 induces cellular senescence in aging muscle through multiple signaling pathways. Aging 6: 160–175.

Hwang HW, Wentzel EA, Mendell JT. 2007. A hexanucleotide element directs microRNA nuclear import. Science 315: 97–100.

Jackson EL, Willis N, Mercer K, Bronson RT, Crowley D, Montoya R, Jacks T, Tuveson DA. 2001. Analysis of lung tumor initiation and progression using conditional expression of oncogenic K-ras. Genes Dev 15: 3243–3248.

Jessen C, Kress JKC, Baluapuri A, Hufnagel A, Schmitz W, Kneitz S, Roth S, Marquardt A, Appenzeller S, Ade CP et al. 2020. The transcription factor NRF2 enhances melanoma malignancy by blocking differentiation and inducing COX2 expression. Oncogene.

Katsuoka F, Yamamoto M. 2016. Small Maf proteins (MafF, MafG, MafK): History, structure and function. Gene 586: 197–205.

Ko T, Sharma R, Li S. 2019. Genome-wide screening identifies novel genes implicated in cellular sensitivity to BRAF(V600E) expression. Oncogene.

Kogure T, Costinean S, Yan I, Braconi C, Croce C, Patel T. 2012. Hepatic miR-29ab1 expression modulates chronic hepatic injury. Journal of cellular and molecular medicine 16: 2647–2654.

Krauthammer M, Kong Y, Bacchiocchi A, Evans P, Pornputtapong N, Wu C, McCusker JP, Ma S, Cheng E, Straub R et al. 2015. Exome sequencing identifies recurrent mutations in NF1 and RASopathy genes in sun-exposed melanomas. Nature genetics 47: 996–1002.

Krauthammer M, Kong Y, Ha BH, Evans P, Bacchiocchi A, McCusker JP, Cheng E, Davis MJ, Goh G, Choi M et al. 2012. Exome sequencing identifies recurrent somatic RAC1 mutations in melanoma. Nature genetics 44: 1006–1014.

Kriegel AJ, Liu Y, Fang Y, Ding X, Liang M. 2012. The miR-29 family: genomics, cell biology, and relevance to renal and cardiovascular injury. Physiological genomics 44: 237–244.

Kunz M, Loffler-Wirth H, Dannemann M, Willscher E, Doose G, Kelso J, Kottek T, Nickel B, Hopp L, Landsberg J et al. 2018. RNA-seq analysis identifies different transcriptomic types and developmental trajectories of primary melanomas. Oncogene 37: 6136–6151.

Kurinna S, Schafer M, Ostano P, Karouzakis E, Chiorino G, Bloch W, Bachmann A, Gay S, Garrod D, Lefort K et al. 2014. A novel Nrf2-miR-29-desmocollin-2 axis regulates desmosome function in keratinocytes. Nat Commun 5: 5099.

Kwon JJ, Factora TD, Dey S, Kota J. 2019. A Systematic Review of miR-29 in Cancer. Molecular therapy oncolytics 12: 173–194.

Liu H, Cheng XH. 2018. MiR-29b reverses oxaliplatin-resistance in colorectal cancer by targeting SIRT1. Oncotarget 9: 12304–12315.

Liu T, Yang H, Fan W, Tu J, Li TWH, Wang J, Shen H, Yang J, Xiong T, Steggerda J et al. 2018. Mechanisms of MAFG Dysregulation in Cholestatic Liver Injury and Development of Liver Cancer. Gastroenterology 155: 557–571 e514.

Love MI, Huber W, Anders S. 2014. Moderated estimation of fold change and dispersion for RNA-seq data with DESeq2. Genome Biology 15: 550.

Lloyd AC, Obermuller F, Staddon S, Barth CF, McMahon M, Land H. 1997. Cooperating oncogenes converge to regulate cyclin/cdk complexes. Genes Dev 11: 663–677.

Martin M. 2011. Cutadapt removes adapter sequences from high-throughput sequencing reads. 2011 **17**: 3 %J EMBnetjournal.

Martinez I, Cazalla D, Almstead LL, Steitz JA, DiMaio D. 2011. miR-29 and miR-30 regulate B-Myb expression during cellular senescence. Proc Natl Acad Sci USA 108: 522–527.

Matsuo M, Nakada C, Tsukamoto Y, Noguchi T, Uchida T, Hijiya N, Matsuura K, Moriyama M. 2013. MiR-29c is downregulated in gastric carcinomas and regulates cell proliferation by targeting RCC2. Mol Cancer 12: 15.

Michaloglou C, Vredeveld LC, Soengas MS, Denoyelle C, Kuilman T, van der Horst CM, Majoor DM, Shay JW, Mooi WJ, Peeper DS. 2005. BRAFE600-associated senescence-like cell cycle arrest of human naevi. Nature 436: 720–724.

Mortazavi A, Williams BA, McCue K, Schaeffer L, Wold B. 2008. Mapping and quantifying mammalian transcriptomes by RNA-Seq. Nat Methods 5: 621–628.

Mott JL, Kobayashi S, Bronk SF, Gores GJ. 2007. mir-29 regulates Mcl-1 protein expression and apoptosis. Oncogene 26: 6133–6140.

Mott JL, Kurita S, Cazanave SC, Bronk SF, Werneburg NW, Fernandez-Zapico ME. 2010. Transcriptional suppression of mir-29b-1/mir-29a promoter by c-Myc, hedgehog, and NF-kappaB. Journal of cellular biochemistry 110: 1155–1164.

Muniyappa MK, Dowling P, Henry M, Meleady P, Doolan P, Gammell P, Clynes M, Barron N. 2009. MiRNA-29a regulates the expression of numerous proteins and reduces the invasiveness and proliferation of human carcinoma cell lines. European journal of cancer 45: 3104–3118.

Muthusamy V, Hobbs C, Nogueira C, Cordon-Cardo C, McKee PH, Chin L, Bosenberg MW. 2006. Amplification of CDK4 and MDM2 in malignant melanoma. Genes, chromosomes & cancer 45: 447–454.

Nan P, Niu Y, Wang X, Li Q. 2019. MiR-29a function as tumor suppressor in cervical cancer by targeting SIRT1 and predict patient prognosis. OncoTargets and therapy 12: 6917–6925.

Nguyen T, Kuo C, Nicholl MB, Sim MS, Turner RR, Morton DL, Hoon DS. 2011. Downregulation of microRNA-29c is associated with hypermethylation of tumor-related genes and disease outcome in cutaneous melanoma. Epigenetics 6: 388­394.

Nishikawa R, Chiyomaru T, Enokida H, Inoguchi S, Ishihara T, Matsushita R, Goto Y, Fukumoto I, Nakagawa M, Seki N. 2015. Tumour-suppressive microRNA-29s directly regulate LOXL2 expression and inhibit cancer cell migration and invasion in renal cell carcinoma. FEBS letters 589: 2136–2145.

Peeper DS, Upton TM, Ladha MH, Neuman E, Zalvide J, Bernards R, DeCaprio JA, Ewen ME. 1997. Ras signalling linked to the cell-cycle machinery by the retinoblastoma protein. Nature 386: 177–181.

Perna D, Karreth FA, Rust AG, Perez-Mancera PA, Rashid M, Iorio F, Alifrangis C, Arends MJ, Bosenberg MW, Bollag G et al. 2015. BRAF inhibitor resistance mediated by the AKT pathway in an oncogenic BRAF mouse melanoma model. Proc Natl Acad SciUSA 112: E536–545.

Romano G, Kwong LN. 2017. miRNAs, Melanoma and Microenvironment: An Intricate Network. Int J Mol Sci 18.

Satyamoorthy K, Li G, Gerrero MR, Brose MS, Volpe P, Weber BL, Van Belle P, Elder DE, Herlyn M. 2003. Constitutive mitogen-activated protein kinase activation in melanoma is mediated by both BRAF mutations and autocrine growth factor stimulation. Cancer Res 63: 756–759.

Schonwasser DC, Marais RM, Marshall CJ, Parker PJ. 1998. Activation of the mitogen-activated protein kinase/extracellular signal-regulated kinase pathway by conventional, novel, and atypical protein kinase C isotypes. Mol Cell Biol 18: 790–798.

Shah NM, Zaitseva L, Bowles KM, MacEwan DJ, Rushworth SA. 2015. NRF2-driven miR-125B1 and miR-29b1 transcriptional regulation controls a novel anti-apoptotic miRNA regulatory network for AML survival. Cell Death Differ 22: 654–664.

Sharpless NE, Kannan K, Xu J, Bosenberg MW, Chin L. 2003. Both products of the mouse Ink4a/Arf locus suppress melanoma formation in vivo. Oncogene 22: 5055–5059.

Sherr CJ, Weber JD. 2000. The ARF/p53 pathway. Current opinion in genetics & development 10: 94–99.

Sheu JJ, Guan B, Tsai FJ, Hsiao EY, Chen CM, Seruca R, Wang TL, Shih Ie M. 2012. Mutant BRAF induces DNA strand breaks, activates DNA damage response pathway, and up-regulates glucose transporter-1 in nontransformed epithelial cells. The American journal of pathology 180: 1179–1188.

Sumimoto H, Imabayashi F, Iwata T, Kawakami Y. 2006. The BRAF-MAPK signaling pathway is essential for cancer-immune evasion in human melanoma cells. The Journal of experimental medicine 203: 1651–1656.

Sun C, Zhang Z, Qie J, Wang Y, Qian J, Wang J, Wu J, Li Q, Bai C, Han B et al. 2018. Genetic polymorphism of SLC31A1 is associated with clinical outcomes of platinum-based chemotherapy in non-small-cell lung cancer patients through modulating microRNA-mediated regulation. Oncotarget 9: 23860–23877.

Teng Y, Zhang Y, Qu K, Yang X, Fu J, Chen W, Li X. 2015. MicroRNA-29B (mir-29b) regulates the Warburg effect in ovarian cancer by targeting AKT2 and AKT3. Oncotarget 6: 40799–40814.

Terzian T, Torchia EC, Dai D, Robinson SE, Murao K, Stiegmann RA, Gonzalez V, Boyle GM, Powell MB, Pollock PM et al. 2010. p53 prevents progression of nevi to melanoma predominantly through cell cycle regulation. Pigment cell & melanoma research 23: 781–794.

Tuveson DA, Shaw AT, Willis NA, Silver DP, Jackson EL, Chang S, Mercer KL, Grochow R, Hock H, Crowley D et al. 2004. Endogenous oncogenic K-ras(G12D) stimulates proliferation and widespread neoplastic and developmental defects. Cancer cell 5: 375–387.

Ugalde AP, Ramsay AJ, de la Rosa J, Varela I, Marino G, Cadinanos J, Lu J, Freije JM, Lopez-Otin C. 2011. Aging and chronic DNA damage response activate a regulatory pathway involving miR-29 and p53. EMBO J 30: 2219–2232.

Vera O, Jimenez J, Pernia O, Rodriguez-Antolin C, Rodriguez C, Sanchez Cabo F, Soto J, Rosas R, Lopez-Magallon S, Esteban Rodriguez I et al. 2017. DNA Methylation of miR-7 is a Mechanism Involved in Platinum Response through MAFG Overexpression in Cancer Cells. Theranostics 7: 4118–4134.

Viros A, Sanchez-Laorden B, Pedersen M, Furney SJ, Rae J, Hogan K, Ejiama S, Girotti MR, Cook M, Dhomen N et al. 2014. Ultraviolet radiation accelerates BRAF-driven melanomagenesis by targeting TP53. Nature 511: 478–482.

Wang H, An X, Yu H, Zhang S, Tang B, Zhang X, Li Z. 2017. MiR-29b/TET1 /ZEB2 signaling axis regulates metastatic properties and epithelial-mesenchymal transition in breast cancer cells. Oncotarget 8: 102119–102133.

Wei W, He HB, Zhang WY, Zhang HX, Bai JB, Liu HZ, Cao JH, Chang KC, Li XY, Zhao SH. 2013. miR-29 targets Akt3 to reduce proliferation and facilitate differentiation of myoblasts in skeletal muscle development. Cell death & disease 4: e668.

Wozniak M, Mielczarek A, Czyz M. 2016. miRNAs in Melanoma: Tumor Suppressors and Oncogenes with Prognostic Potential. Current medicinal chemistry 23: 3136–3153.

Wu GS. 2004. The functional interactions between the p53 and MAPK signaling pathways. Cancer biology & therapy 3: 156–161.

Wu J, Rosenbaum E, Begum S, Westra WH. 2007. Distribution of BRAF T1799A(V600E) mutations across various types of benign nevi: implications for melanocytic tumorigenesis. The American Journal of dermatopathology 29: 534–537.

Wu Z, Huang X, Huang X, Zou Q, Guo Y. 2013. The inhibitory role of Mir-29 in growth of breast cancer cells./ Exp Clin Cancer Res 32: 98.

Xiong Y, Liu L, Qiu Y, Liu L. 2018. MicroRNA-29a Inhibits Growth, Migration and Invasion of Melanoma A375 Cells in Vitro by Directly Targeting BMI1. Cellular physiology and biochemistry: international journal of experimental cellular physiology, biochemistry, and pharmacology 50: 385–397.

Yeh I, von Deimling A, Bastian BC. 2013. Clonal BRAF mutations in melanocytic nevi and initiating role of BRAF in melanocytic neoplasia. Journal of the National Cancer Institute 105: 917–919.

Yu H, McDaid R, Lee J, Possik P, Li L, Kumar SM, Elder DE, Van Belle P, Gimotty P, Guerra M et al. 2009. The role of BRAF mutation and p53 inactivation during transformation of a subpopulation of primary human melanocytes. The American journal of pathology 174: 2367–2377.

Yu PN, Yan MD, Lai HC, Huang RL, Chou YC, Lin WC, Yeh LT, Lin YW. 2014. Downregulation of miR-29 contributes to cisplatin resistance of ovarian cancer cells. International journal of cancer 134: 542–551.

Yuspa SH, Harris CC. 1974. Altered differentiation of mouse epidermal cells treated with retinyl acetate in vitro. Experimental cell research 86: 95–105.

Zhang T, Dutton-Regester K, Brown KM, Hayward NK. 2016. The genomic landscape of cutaneous melanoma. Pigment cell & melanoma research 29: 266–283.

Zhang X, Zhao X, Fiskus W, Lin J, Lwin T, Rao R, Zhang Y, Chan JC, Fu K, Marquez VE et al. 2012. Coordinated silencing of MYC-mediated miR-29 by HDAC3 and EZH2 as a therapeutic target of histone modification in aggressive B-Cell lymphomas. Cancer cell 22: 506–523.

Zhang Y, Yang L, Wang S, Liu Z, Xiu M. 2018. MiR-29a suppresses cell proliferation by targeting SIRT1 in hepatocellular carcinoma. Cancer biomarkers: section A of Disease markers 22: 151–159.

Zhao JJ, Lin J, Lwin T, Yang H, Guo J, Kong W, Dessureault S, Moscinski LC, Rezania D, Dalton WS et al. 2010. microRNA expression profile and identification of miR-29 as a prognostic marker and pathogenetic factor by targeting CDK6 in mantle cell lymphoma. Blood 115: 2630–2639.

