## Supplementary Figures and Legends for "MAPK-induced miR-29 targets MAFG and suppresses melanoma development"

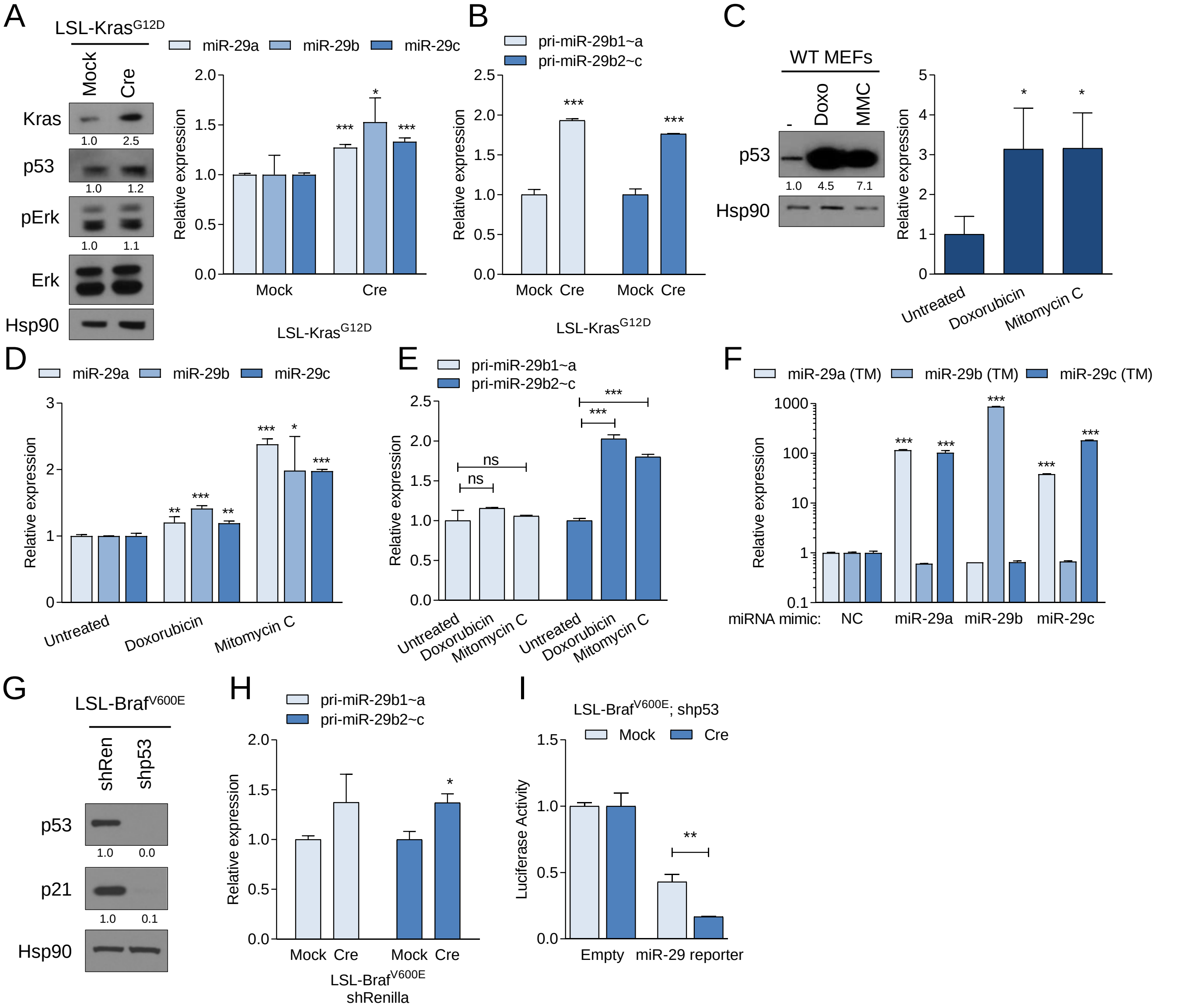


**Supplementary Figure 1:** (A) Induction of Kras$^{G12D}$ expression by Adeno-Cre (Cre) in LSL-Kras$^{G12D}$ MEFs elevates expression of p53 (Western blot, left panel) and mature miR-29a, miR-29b, and miR-29c (qRT-PCR, right panel). MEFs infected with empty adenovirus (Mock) serve as controls. (B) qRT-PCR showing the expression of pri-miR-29b1~a and pri-miR-29b2~c in Kras$^{G12D}$ MEFs following Adeno-Cre/Mock infection. (C) Effect of Doxorubicin (Doxo) and Mitomycin C (MMC) on p53 (left) and p21 (right) in wiltdype MEFs. (D) Quantitative expression of the mature form of miR-29a, -29b and -29c after induction of p53-dependent stress by Doxorubicin and Mitomycin C measured by qRT-PCR. (E) qRT-PCRs showing the expression of pri-miR-29b1~a and pri-miR-29b2~c in wildtype MEFs following treatment with Doxorubicin or Mitomycin C. (F) Quantification by qRT-PCR of mature of miR-29a, -29b and -29c after overexpression of microRNA mimics in A375 cells. Cells overexpressing a negative control mimic were used as controls. TM, TaqMan probe. (G) Confirmation of p53 silencing in LSL- Braf$^{V600E}$ MEFs infected with shp53 retrovirus. (H) Expression of pri-miR-29b1~a and pri-miR-29b2~c measured by qRT-PCR after Cre-mediated activation of Braf$^{V600E}$ in shRenilla LSL-Braf$^{V600E}$ MEFs. (I) Luciferase assay in shp53-expressing LSL-Braf$^{V600E}$ MEFs using a miR-29-Luciferase reporter. For Western blots, Hsp90 was used as loading control. The mean ± SEM of one representative out of two independent experiments performed in triplicate is shown. Gene expression levels are normalized to β-Actin as endogenous control. ns, not significant; * $p < 0.05$; ** $p < 0.01$; *** $p < 0.001$.

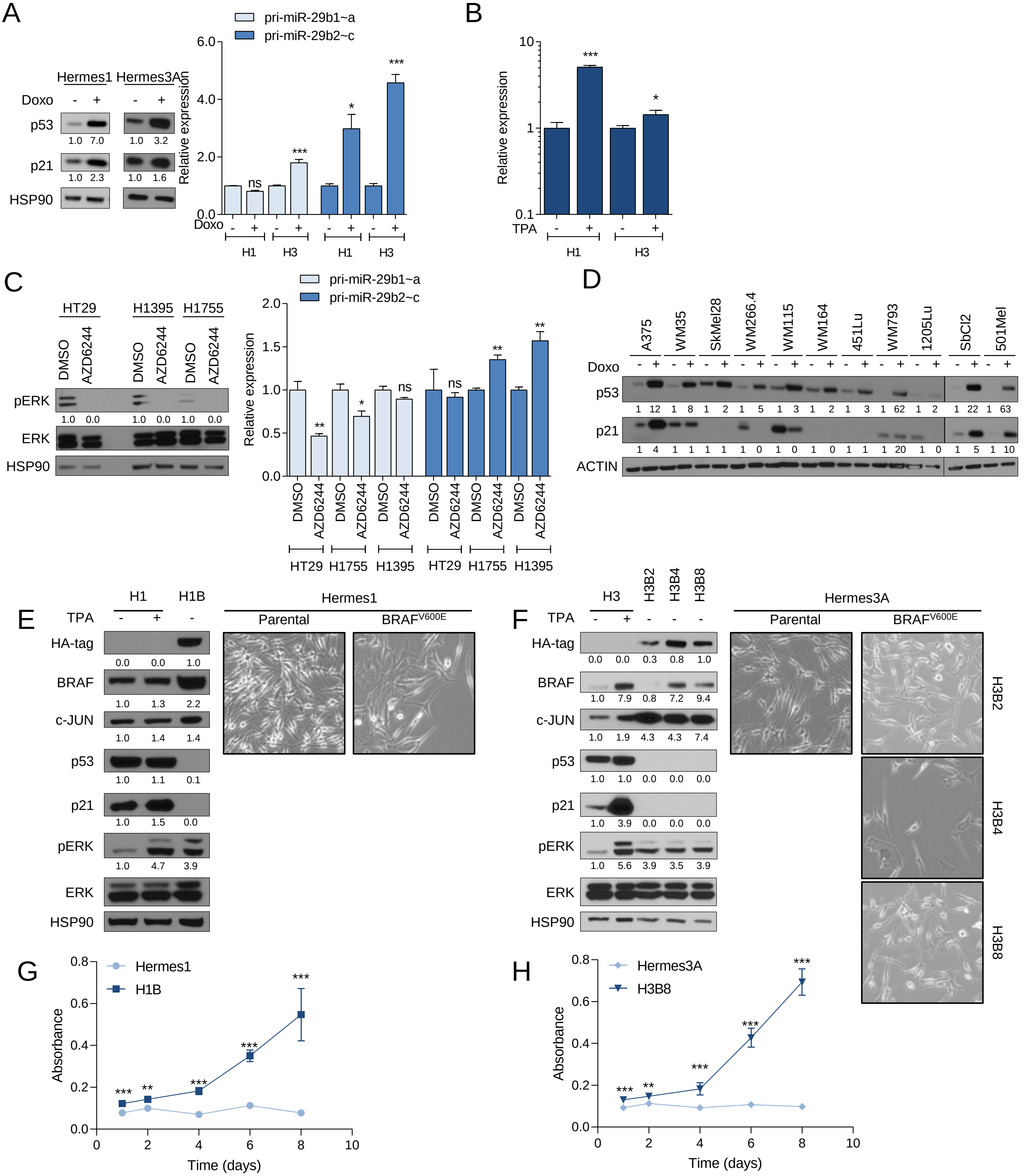

**Supplementary Figure 2:** (A) Effect of Doxorubicin (Doxo) on pri-miR-29b1~a and pri-miR-29b2~c expression in human melanocytes Hermes1 (H1) and Hermes3A (H3). (B) Effect of TPA on the MAPK downstream effector c-JUN in human melanocytes Hermes1 (H1) and Hermes3A (H3). (C) Effect of AZD6244 on primiR-29b1~a and pri-miR-29b2~c expression in BRAF mutant colon (HT29) and lung (H1395, H1755) cancer cell lines. (D) Effect of Doxorubicin (Doxo) on p53 and p21 in a panel of BRAF and NRAS mutant melanoma cell lines. (E-F) Comparison of protein expression and morphology between parental melanocytes Hermes1 (H1, E) and Hermes3A (H3, F) cultured with (+) and without TPA (-) and BRAF$^{V600E}$ melanocytes (H1B, H3Bs) in absence of TPA (-). (G-H) Proliferation assay comparing Hermes1 with H1B (G) and Hermes3A with H3B8 (H) cell lines cultured in absence of TPA. The combined mean ± SEM of two independent experiments performed in quadruplicate is shown. Ns, not significant; * $p < 0.05$; ** $p < 0.01$; *** $p < 0.001$.

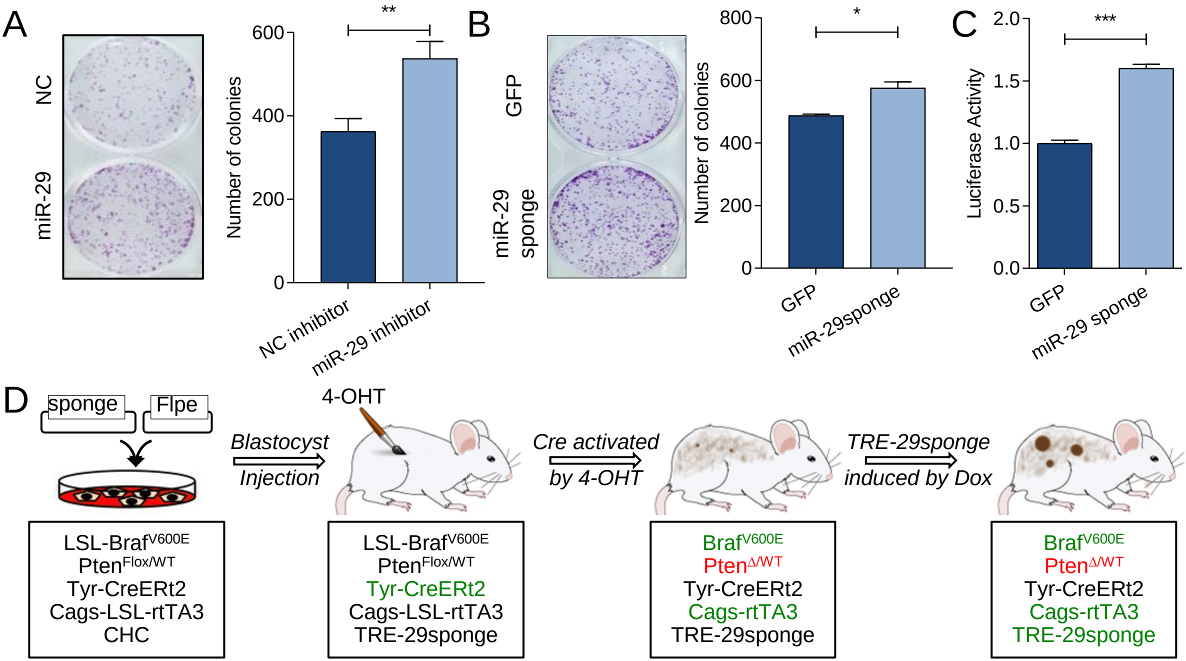


**Supplementary Figure 3.** (A) Effect of miR29 inactivation by hairpin inhibitors on colony formation of A375 cells . (B) Effect of miR-29 inactivation by a miR-29 sponge construct on colony formation of A375 cells. The mean ± SEM of one representative out of two independent experiments performed in triplicate is shown in (A) and (B). (C) Effect of the miR-29 sponge on miR-29-Luciferase reporter activity in A375 cells. The combined mean ± SEM of three independent experiments performed in quadruplicate is shown. (D) Outline of the embryonic stem cell-genetically engineered mouse model approach (Bok et al. 2019) where a Dox-inducible miR-29 sponge is expressed in $Braf^{V600E}$; $Pten^{\Delta/WT}$ melanocytes. * $p < 0.05$; ** $p < 0.01$; *** $p < 0.001$

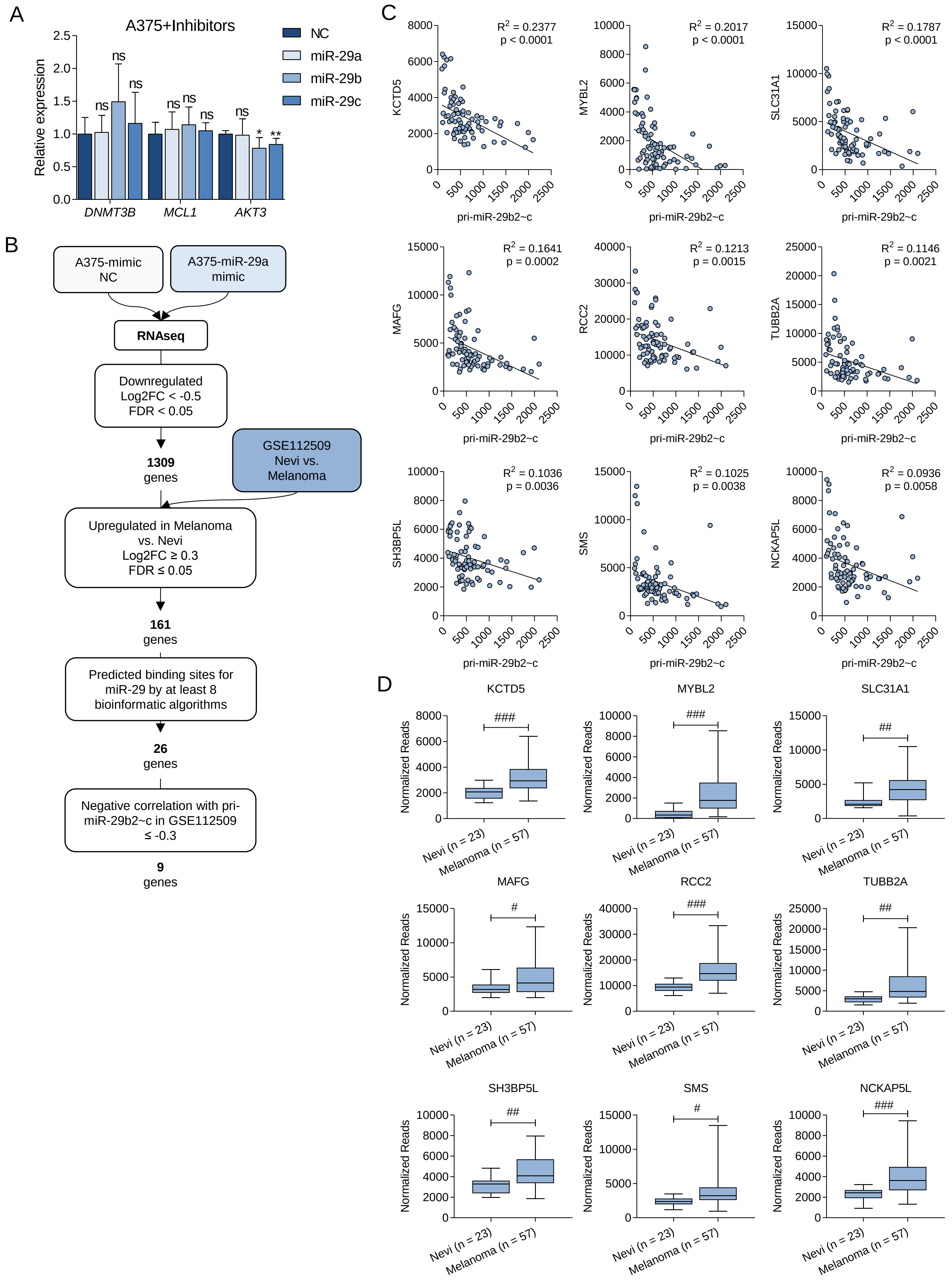


**Supplementary Figure 4.** (A) Quantification by qRT-PCR of three validated miR-29 target genes of miR-29 after overexpression of miR-29 mimics in A375 cells. Cells overexpressing a negative control mimic were used as controls. (B) Selection flowchart to identify target genes of miR-29 involved in melanoma progression. (C) Correlation between pri-miR-29b2~c and the top 9 identified putative miR-29 targets in the GSE112509 dataset. (D) Expression of the top 9 identified putative miR-29 targets in 23 nevi and 57 melanomas obtained from the GSE112509 dataset. # FDR < 0.05; ## FDR < 0.01; ### FDR < 0.001.

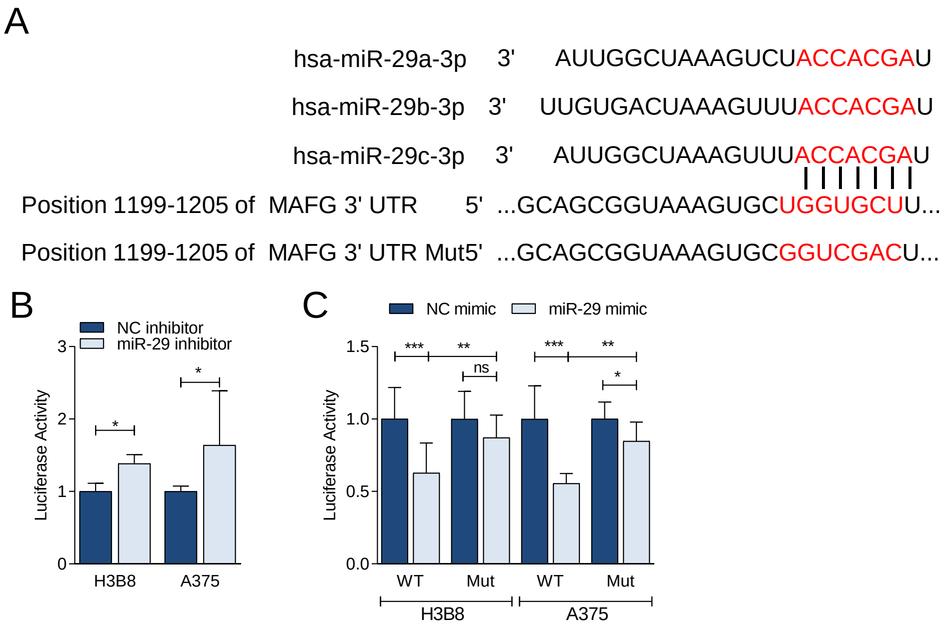


**Supplementary Figure 5.** (A) Alignment of mature miR-29a, -29b and -29c with the miR-29 binding site in the human MAFG 3'UTR. The mutates miR-29 binding site used in the Luciferase reporter assays is shown below. (B) Activity of 3'UTR Luciferase reporter in response to miR-29 inhibitors. (C) Activity of MAFG wildtype or miR-29 binding site-mutant MAFG 3'UTR Luciferase reporter in response to miR-29 mimics. The combined mean ± SEM of two independent experiments performed in quadruplicate is shown. Ns, not significant; * $p < 0.05$; ** $p < 0.01$; *** $p < 0.001$.

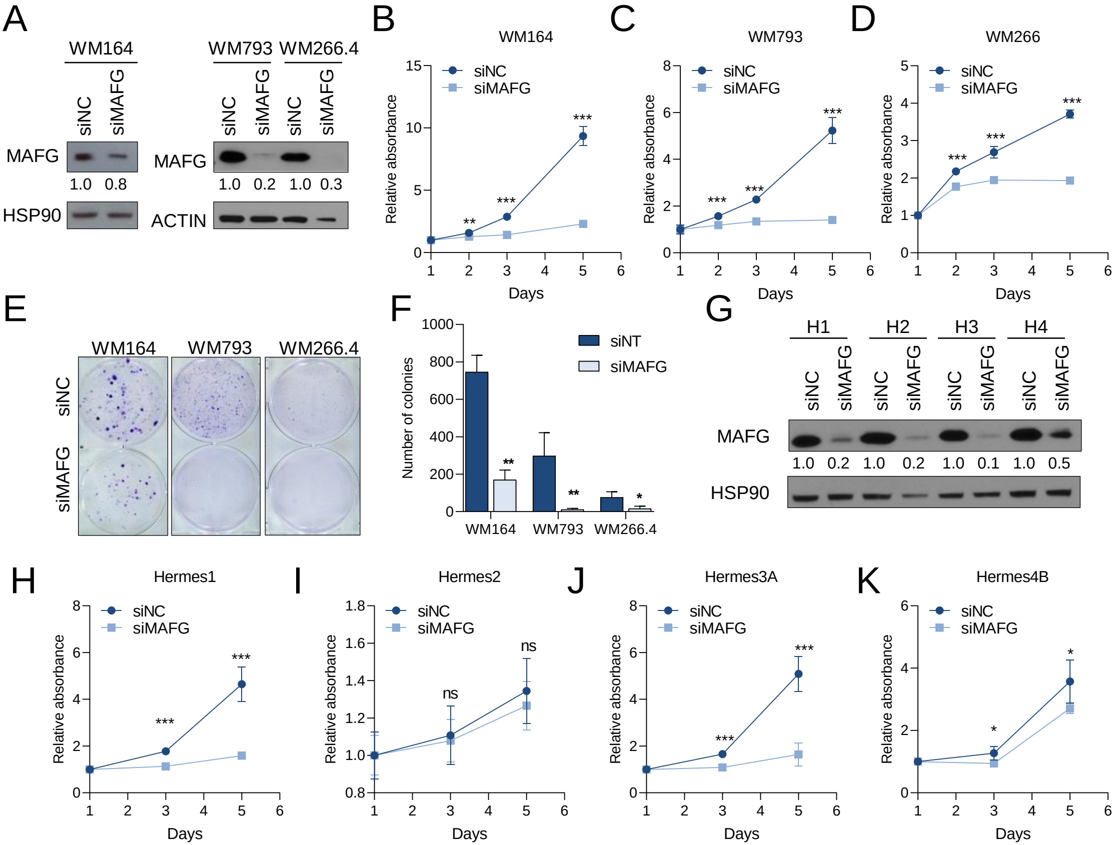


**Supplementary Figure 6**. (A) Silencing of MAFG in melanoma cells WM164, WM793 and WM266.4 with an ON-TARGETplus MAFG siRNA pool. (B-F) Proliferation (B-D) and colony formation (E,F) of WM164 (B,E), WM793 (C,E) or WM266.4 (D,E) cells upon MAFG silencing. (G) Silencing of MAFG in Hermes1-4 melanocytes with an ON-TARGETplus MAFG siRNA pool. (H-K) Effect of MAFG silencing on proliferation of Hermes1 (H), Hermes2 (I), Hermes3A (J) and Hermes4B (K). For Western blots, HSP90 or β-ACTIN were used as loading controls. Ns, not significant; * $p < 0.05$; ** $p < 0.01$; *** $p < 0.001$.

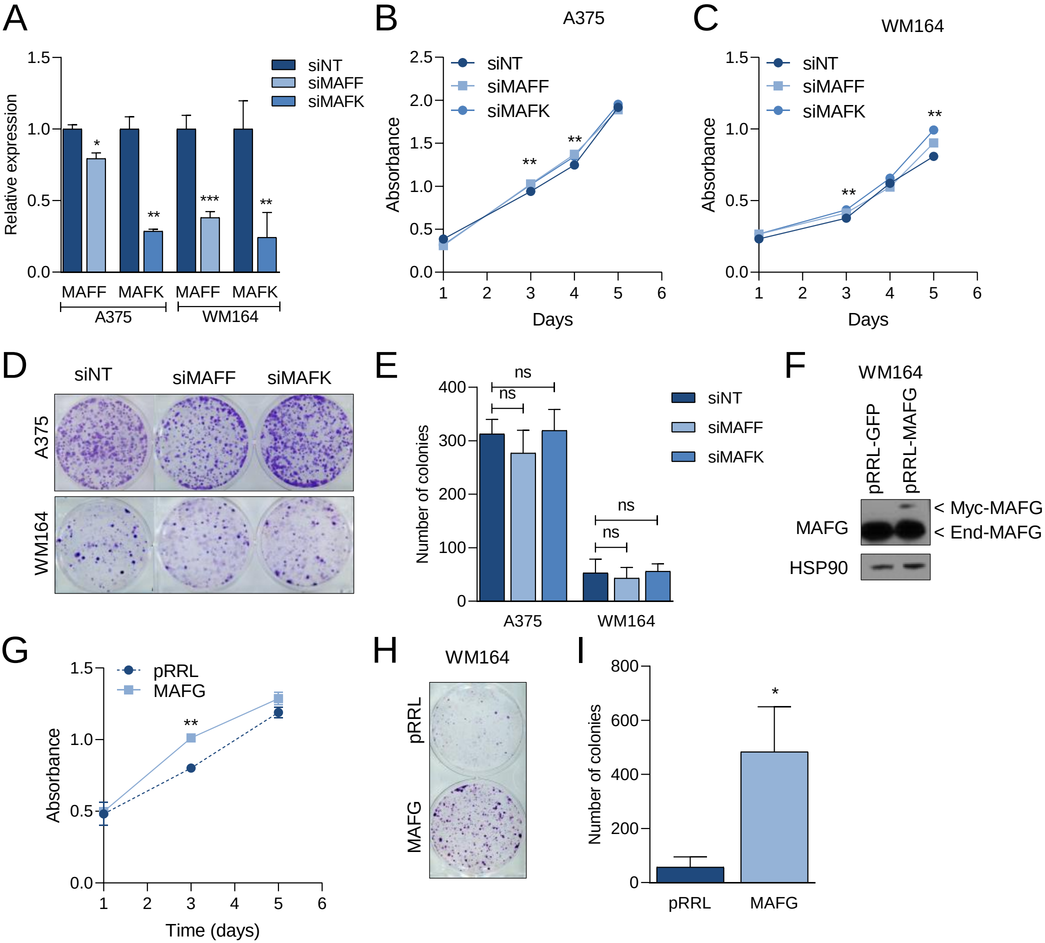


**Supplementary Figure 7**. (A) qRT-PCR validation of the silencing of MAFF and MAFK in melanoma cells A375 and WM164 with an ON-TARGETplus MAFF or MAFK siRNA pool. (B-E) Proliferation (B, C) and colony formation (D,F) of A375 (B,D) or WM164 (C, D) cells upon MAFF and MAFK silencing. (F) Overexpression of MAFG in WM164 cells with a Dox-inducible vector. (G-I) Effect of MAFG overepxression on proliferation (G) and focus formation (H,I) of WM164. For Western blots, HSP90 was used as loading control. ns, not significant; * $p < 0.05$; ** $p < 0.01$; *** $p < 0.001$.
